# Non-parametric modelling of temporal and spatial counts data from RNA-seq experiments

**DOI:** 10.1101/2020.07.29.227207

**Authors:** Nuha BinTayyash, Sokratia Georgaka, ST John, Sumon Ahmed, Alexis Boukouvalas, James Hensman, Magnus Rattray

## Abstract

**Motivation:** The negative binomial distribution has been shown to be a good model for counts data from both bulk and single-cell RNA-sequencing (RNA-seq). Gaussian process (GP) regression provides a useful non-parametric approach for modeling temporal or spatial changes in gene expression. However, currently available GP regression methods that implement negative binomial likelihood models do not scale to the increasingly large datasets being produced by single-cell and spatial transcriptomics.

**Results:** The GPcounts package implements GP regression methods for modelling counts data using a negative binomial likelihood function. Computational efficiency is achieved through the use of variational Bayesian inference. The GP function models changes in the mean of the negative binomial likelihood through a logarithmic link function and the dispersion parameter is fitted by maximum likelihood. We validate the method on simulated time course data, showing that it is better able to identify changes in over-dispersed counts data than methods based on Gaussian or Poisson likelihoods. To demonstrate temporal inference, we apply GPcounts to single-cell RNA-seq datasets after pseudotime and branching inference. To demonstrate spatial inference, we apply GPcounts to data from the mouse olfactory bulb to identify spatially variable genes and compare to two published GP methods. We also provide the option of modelling additional dropout using a zero-inflated negative binomial. Our results show that GPcounts can be used to model temporal and spatial counts data in cases where simpler Gaussian and Poisson likelihoods are unrealistic.

**Availability:** GPcounts is implemented using the GPflow library in Python and is available at https://github.com/ManchesterBioinference/GPcounts along with the data, code and notebooks required to reproduce the results presented here.

## 1 Introduction

Biological data are often summarised as counts, e.g. high-throughput sequencing allows us to count the number of sequence reads aligning to a genomic region of interest, while single molecule fluorescence in-situ hybridization (smFISH) allows us to count the number of RNA molecules in a cell or within a cellular compartment. For datasets that are collected with temporal or spatial resolution, it can be useful to model changes in time or space using a Gaussian process (GP). GPs provide a flexible framework for non-parametric Bayesian modelling, allowing for non-linear regression while quantifying the uncertainty associated with the estimated latent function and data measurement process (Rasmussen and Williams, 2006). GPs are useful methods for analysing gene expression time series data, e.g. to identify differentially expressed genes, model temporal changes across conditions, cluster genes or model branching dynamics (Stegle *et al.*, 2010; Kalaitzis and Lawrence, 2011; Äijö *et al.*, 2014; Yang *et al.*, 2016; Ahmed *et al.*, 2019; Hensman *et al.*, 2013; McDowell *et al.*, 2018; Boukouvalas *et al.*, 2018). GPs have also been applied to spatial gene expression data as a method to discover spatially varying genes (Svensson *et al.*, 2018; Sun *et al.*, 2020). In many previous applications of GP regression to counts, the log-transformed counts data are modelled using a Gaussian noise assumption. Such Gaussian likelihood models do not properly capture the typical characteristics of counts data, in particular substantial probability mass at zero counts and heteroscedastic noise. Alternative data transformations attempt to match the mean/variance relationship of counts data (Anscombe, 1948) but cannot model all relevant aspects of the distribution, e.g. the probability mass at zero. More suitable distributions for modelling counts data include the negative binomial distribution, which can model over-dispersion beyond a Poisson model, and zero-inflated extensions that have been developed for single-cell data exhibiting an excess of zero counts (Pierson and Yau, 2015; Risso *et al.*, 2018). A negative binomial likelihood was implemented for GP regression using Markov chain Monte Carlo (MCMC) inference (Äijö *et al.*, 2014). However, this approach does not scale to large datasets, e.g. single-cell RNA-seq (scRNA-seq) data after pseudotime inference or from spatial transcriptomics assays.

Various statistical methods have been proposed for spatially resolved omics data (Svensson *et al.*, 2018; Sun *et al.*, 2020; Arnol *et al.*, 2019; Edsgärd *et al.*, 2018). The SpatialDE approach uses a GP to model the expression variability of each gene with a spatial and non-spatial component (Svensson *et al.*, 2018). The latter is modelled as observation noise, while the former is modelled using a covariance function that depends on the pairwise distance between the cells. The ratio of these components is used to measure the spatial variability of each gene. A Gaussian likelihood is used for inference and the raw counts data is transformed using Anscombe’s transformation (Anscombe, 1948) to reduce heteroscedasticity. Statistical significance is assessed by a likelihood ratio test against a spatially homogeneous model, with p-values calculated under the assumption of a χ^2^-distribution. Trendsceek (Edsgärd *et al.*, 2018) assesses spatial expression of each gene by modelling normalised counts data as a marked point process, where the points represent the spatial locations and marks represent the expression levels, with a permuted null used to assess significance. SPARK (Sun *et al.*, 2020) is a recently introduced GP-based method that relies on a variety of spatial kernels and a penalized quasi-likelihood algorithm. SPARK models counts using a Poisson likelihood function and captures over-dispersion through a nugget (white noise) term in the underlying GP covariance function.

Here, we introduce an alternative GP regression model (GPcounts) to model temporal or spatial counts data with a negative binomial (NB) likelihood. GPcounts can be used for a variety of tasks, e.g. to infer temporal trajectories, identify differentially expressed genes using one-sample or two-sample tests, infer branching genes from scRNA-seq after pseudotime inference, or to infer spatially varying genes. We use a GP with a logarithmic link function to model variation in the mean of the counts data distribution across time or space. As an example, in Figure 1 we show how GPcounts captures the distribution of a short RNA-seq time course dataset from Leong *et al.* (2014). In this example we use a two-sample test to determine whether time-series measured under different conditions have differing trajectories. The GP with Gaussian likelihood fails to model the data distribution well, leading to a less plausible inferred dynamics and overly broad credible regions. Although the trajectories of the two samples clearly differ, the Gaussian likelihood model provides a poor fit and the single trajectory model is preferred in the two-sample test.

**Figure 1:**
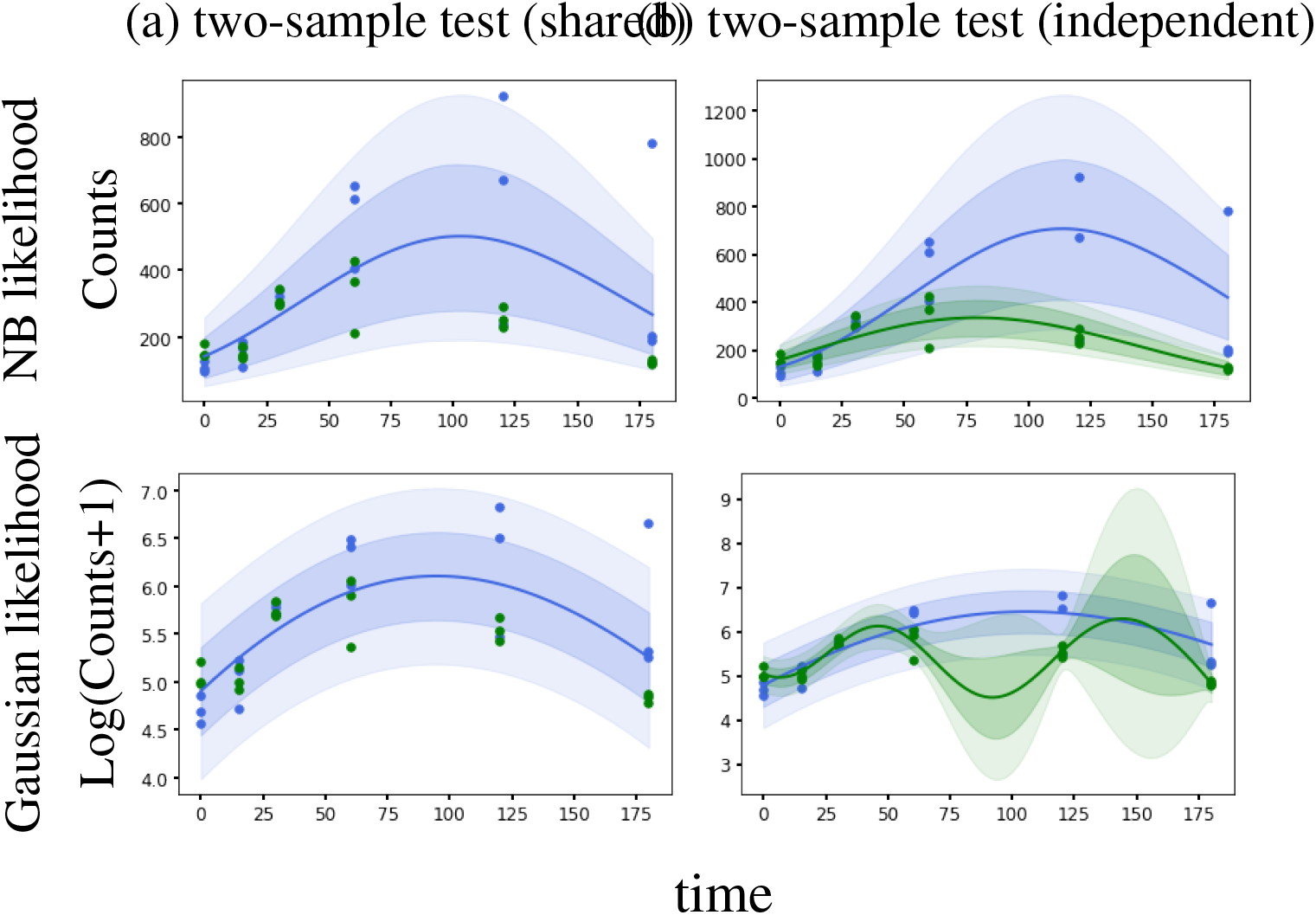
Example of data from a two-sample time course RNA-seq experiment (Leong *et al.*, 2014). With a Negative Binomial likelihood we are able to identify this differentially expressed gene based on a likelihood ratio statistic. With a Gaussian noise model we obtain a poor fit and the likelihood ratio does not identify the gene as differentially expressed.

Our package is developed using the GPflow library (De G. Matthews *et al.*, 2017) which we have extended to include the DEtime kernel (Yang *et al.*, 2016) for branching genes and to include an NB likelihood function with an efficient variational approximate inference method.

## 2 Methods

### 2.1 Gaussian Process Regression

A Gaussian Process (GP) is a stochastic process over real valued functions and defines a probability distribution over function space (Rasmussen and Williams, 2006). GPs provide a non-linear and non-parametric framework for inference. Consider a set of *N* temporal or spatial locations ***x*** = [*x*_1_, *x*_2_,…, *x_N_*] associated with observations ***y*** = [*y*_1_, *y*_2_,…, *y_N_*].

As this is a continuous model, spacings between data points can vary. Each observation *y_n_* is modelled as a noisy observation of the function evaluated at *x_n_*, *f_n_* = *f* (*x_n_*), through some likelihood function *p*(*y_n_*|*f_n_*). The function *f* is a latent function sampled from a GP prior and*p*(*y_n_*|*f_n_*) models the data distribution at *x* = *x_n_*. The simplest and most popular choice of likelihood function is i.i.d. Gaussian noise centred at *f*, in which case 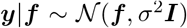, where **f** = [*f*_1_, *f*_2_,…, *f_N_*] is the GP function evaluated at the times or locations in ***x***. In this work we use likelihood functions that are more suitable for counts data, which we introduce below.

To indicate that *f* is drawn from a GP we write,

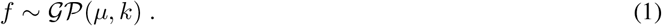

Here, *μ*(*x*) = E[*f*(*x*)] is the mean function of the GP. The covariance function *k* (also known as the kernel function) is a positive semidefinite function 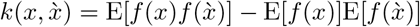 that determines the covariance of *f* at any two locations *x* and 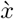. The covariance function describes our beliefs when modelling the function, e.g. whether it is stationary or non-stationary, rough or smooth etc. A popular choice is the Radial Basis Function (RBF) kernel which leads to smooth and infinitely differentiable functions, defined as:

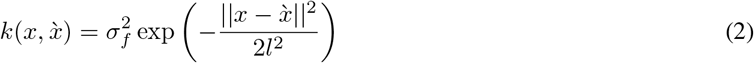

where the hyper-parameters are the lengthscale *l*, controlling spatial or temporal variability, and the amplitude 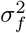, controlling the marginal variance of the function. The RBF kernel is also known as the squared exponential, exponentiated quadratic or Gaussian kernel (Rasmussen and Williams, 2006). Below we describe an alternative kernel appropriate for branching functions.

The kernel hyper-parameters and parameters of the likelihood function can be learned from the data by optimising the log marginal likelihood function of the GP. The marginal likelihood is given by the probability of the data **y** after integrating out the GP function,

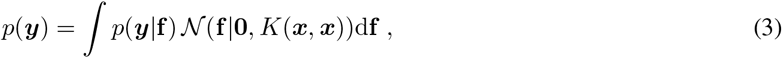

where we have set the GP prior mean to be zero and *K*(***x***, ***x***) is the covariance matrix of the GP function evaluated at the data locations ***x***. For the special case of a Gaussian likelihood function this integral is tractable but for the non-Gaussian likelihoods below we use a variational approximation introduced by Opper and Archambeau (2009).

GPcounts enables model comparison between models with alternative kernels, such as linear and periodic. These kernels are defined as:

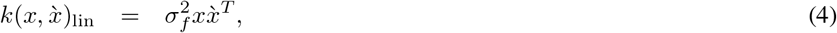

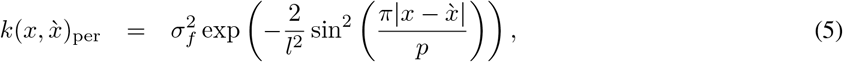

where *p* is the period. When comparing different covariance functions the models are assesed based on the Bayesian Information Criterion (BIC), defined as:

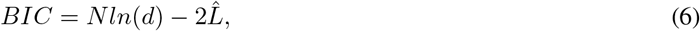

where 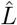 is the log marginal likelihood, *N* corresponds to the number of observations and *d* to the number of optimised hyper-parameters of a given model.

### 2.2 Negative binomial likelihood

The Negative Binomial (NB) distribution is a popular model for bulk RNA-seq counts data (Robinson and Smyth, 2007; Love *et al.*, 2014) and has been shown to be suitable for modelling scRNA-seq count data with UMI (unique molecular identifier) normalisation (Svensson, 2020; Townes *et al.*, 2019). It can be parametrised by a fixed number of failures *r* and mean *μ*:

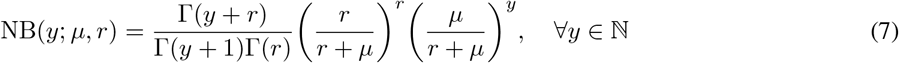

where Γ is the Gamma function. It is convenient to parametrise the NB distribution by a dispersion parameter *α* = *r*^-1^ which captures excess variance relative to a Poisson distribution, since Var[*y*] = *μ* + *αμ*^2^. A logarithmic link function is used to model the mean of the NB as a transformation of the GP function *f*(*x*) = log *μ*(*x*).

In some cases it has been found useful to model additional zero counts through a zero-inflated negative binomial (ZINB) distribution (Pierson and Yau, 2015; Risso *et al.*, 2018). Such extensions can easily be implemented by modifying the likelihood function and we provide a specific implementation using a Michaelis-Menten drop-out function (see Supplementary Material). However, for modern UMI-normalised datasets the standard NB likelihood is often sufficient for inference and estimating an additional drop-out parameter can be difficult (Choi *et al.*, 2020).

### 2.3 Tests and credible regions

We provide one-sample and two-sample likelihood-ratio statistics in GPcounts to identify differentially expressed genes. In the one-sample case the null hypothesis assumes there is no difference in gene expression across time or space (Kalaitzis and Lawrence, 2011; Svensson *et al.*, 2018) and we compute the ratio of the GP model marginal likelihood versus a constant model likelihood. The models are nested since the constant model is equivalent to the GP with an infinite lengthscale parameter. In the two-sample case the null hypothesis assumes there is no difference in gene expression under two different conditions and the alternative hypothesis is that two different GP functions are required to model each sample (Stegle *et al.*, 2010). Rejecting the null hypothesis in each case indicates that a gene is differentially expressed. In the two-sample case, we fit three GPs: one for each dataset separately and a shared GP where the datasets are treated as replicates.

For spatial transcriptomics data we follow two testing procedures. The approach of Svensson *et al.* (2018) is to use *p*-values estimated according to a *χ*^2^-distribution and a 5% FDR threshold is estimated using the approach of Storey and Tibshirani (2003). To obtain better FDR calibration we also implement a permutation test where we randomly rearrange the spatial coordinates to estimate *p*-values based on the permuted null.

To plot [5%-95%] credible regions we draw 100 random samples from the GP at 100 equally spaced times. We exponentiate each GP sample to set the mean of the count distribution (NB or Poisson) and draw counts at each time. We use the mean and percentiles to plot the predictive distribution with the associated credible regions. To smooth the mean samples, we use the Savitzky-Golay filter with cubic polynomial to fit the samples (Savitzky and Golay, 1964). To smooth the samples of the credible regions, we use the Locally Weighted Scatterplot Smoothing (LOWESS) method (Cleveland, 1979).

### 2.4 Scale normalisation for gene expression data

In some cases there may be confounding variation that will dominate the temporal or spatial trends in the data. For example, Svensson *et al.* (2018) point out that there may be spatial patterns in cell size that can lead to almost all genes being identified as spatially variable. In this case it is necessary to normalise away such confounding variation in order to model other sources of spatial variation. Svensson *et al.* (2018) used ordinary least-squares regression of Anscombe-normalised spatial expression data against logged total counts to remove this confounding variation. We use NB regression with an identity link function to learn location-specific normalisation factors for spatial counts data. In GPcounts, we model the temporal or spatial data using a modified GP likelihood *y_i_* ~ NB(*k_i_μ_i_*, *r*) for *i* = 1,…, *N* with *μ_i_* = *e*^*f*(*x_i_*)^ and 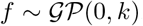. In spatial data, the multiplicative normalisation factor *k_i_* is calculated as *k_i_* = *βT_i_* for *i* = 1,…, *N*, where *β* is the slope calculated by fitting for each gene a NB regression model with intercept zero and *T_i_* corresponds to the total counts at the *i*th spatial location.

### 2.5 Modelling branching dynamics

In the two-sample time course setting it can also be useful to identify the time at which individual genes begin to follow different trajectories. This can be useful in bulk time course data in a two-sample testing scenario (Yang *et al.*, 2016) or for modelling branching dynamics in single-cell data after pseudotime inference (Boukouvalas *et al.*, 2018). We first define a joint covariance function for two GPs with the same covariance function 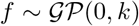 and 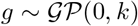 constrained to cross at a branching point *x_b_*:

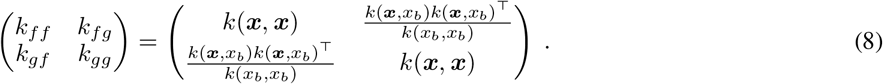

Now consider data from two lineages ***y***^*t*^ and ***y***^*b*^ representing noise-corrupted measurements of a baseline (trunk) and diverging (branch) time course respectively. Before the divergence point *x_b_* the data are distributed around the trunk function *f*(*x*) according to some likelihood function *p*(*y*|*f*),

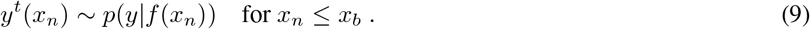

After the branching point *x_b_*, the mean function of the trunk continues to follow *f* while the mean function of the diverging branch trajectory changes to follow *g*,

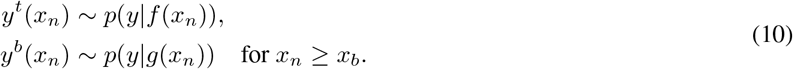

The branching point *x_b_* is a hyper-parameter of the joint covariance function of this model along with the hyper-parameters of the GP functions (lengthscale *l* and amplitude 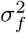). The lengthscale and amplitude are shared by both functions and are fitted (along with the likelihood model parameters) by fitting two separate GP regression models to the data from both conditions or lineages and estimated using maximum likelihood (Yang *et al.*, 2016). This leaves the problem of inferring the branching time *x_b_* only. As this is a one-dimensional problem, the posterior distribution of *x_b_* is estimated using a simple histogram approach. We have used a simple discretisation *x_b_* ∈ [*x*_min_, *x*_min_ + *δ*, *x*_min_ + 2δ,…, *x_max_*] as in Yang *et al.* (2016) and estimate the posterior by using the normalised likelihood evaluated at each grid point,

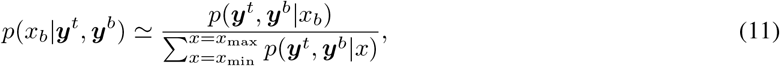

which avoids the need for complex optimisation or integration schemes.

### 2.6 Efficient approximate inference implementation

The integral in Eqn. (3) is intractable for non-Gaussian likelihood functions. Therefore, we need to approximate the posterior and variational inference provides a computationally efficient approximation (Bernardo *et al.*, 2003; Opper and Archambeau, 2009; Bauer *et al.*, 2016). Full variational Bayesian inference is computationally intensive and requires 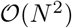 memory and 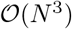 computation time so we use a sparse approximation to reduce the computational requirements (Seeger, 2000; Rasmussen and Williams, 2006). In sparse inference, we choose *M* < *N* inducing points *z* defined in the same space of regressors ***x***. Using inducing points reduces the time complexity to 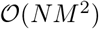. In GPcounts the default is to set the number of inducing points to *M* = 5%(*N*) although less should be used for very large datasets. We apply the *ϕ*-approximate M-determinantal point process (M-DPP) algorithm to select the candidate inducing points (Burt *et al.*, 2020). Using the M-DPP algorithm reduces the required number of inducing points to 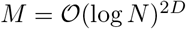 in the case of a squared exponential kernel, where *D* is the data dimensionality. Therefore, a smaller number of inducing points can be used for very large datasets and we provide the user with the option to set the number of inducing points. We also provide the option to use a simpler k-means clustering algorithm to select the inducing point locations (Hensman *et al.*, 2015).

### 2.7 Practical considerations

Practical limitations of the inference framework include local optima in the hyperparameter search and occasional numerical instabilities. We have therefore incorporated some checks to detect both numerical errors failure of Cholesky decomposition or failure of optimization. Local optima are a likely issue if the GP posterior predictive is consistently higher or lower than the observations. Where we suspect local optima or numerical errors we restart the optimisation from new random values for the hyper-parameters. Figure 1 in the Supplementary Material shows an example from a two-sample time course RNA-seq experiment (Leong *et al.*, 2014) with a NB likelihood, where the GP is facing a local optimum solution and safe mode option is switched off in (a) while in (b) safe mode option is switched on so that GPcounts can detect and fix the problem.

## 3 Results and discussion

Below we apply GPcounts on simulated counts time course datasets, scRNA-seq data after pseudotime inference and a spatial transcriptomics dataset.

### 3.1 Assessment on synthetic time course data

We simulated four counts time course datasets with two levels of mean expression (high/low) and two levels of dispersion (high/low) to assess the performance of GPcounts in identifying differentially expressed genes using a one-sample test. Each dataset has 600 genes with time-series measurements at 11 equally spaced time points *x* = [0,0.1,0.2,… 1.0] and two replicates at each time. Half of the genes are differentially expressed across time. We use two classes of generative functions *f* (*x*), sine functions and cubic splines, to simulate data from time-varying genes. The sine functions are of the form *f* (*x*) = *a* sin(*xb* + *d*) + *c* where parameters are drawn from uniform distributions. Specifically, *b* ~ *U*[*π*/4,2*π*], *d* ~ *U*[0, 2*π*] while the *a* and *c* distributions are chosen to alter the signal amplitude and mean ranges respectively. The cubic spline function has the form *f* (*x*) ∈ *C*^2^ [0,1] passing through two control points (*x, y*) where *x* and *y* are drawn from uniform distributions. For non-differentially expressed genes we choose constant values set to the median value of each dynamic function in the dataset. The low and high dispersion values are drawn from uniform distributions *α*_low_ ~ *U*[0.01,0.1] and *α*_high_, ~ *U*[1,3] respectively. An exponential inverse-link function is used to determine the mean of count data at each time *μ*(*x*) = *e*^*f*(*x*)^ and we use the SciPy library (Millman and Aivazis, 2011) to sample counts from the negative binomial distribution. Specific simulation parameters and final datasets are provided with the supporting code.

In Figure 2 we compare the performance of GPcounts using the NB likelihood with a Poisson likelihood and the Gaussian likelihood using a simple logarithmic transformation log(*y* + 1) or using Anscombe’s transformation (Anscombe, 1948) as implemented in SpatialDE (Svensson *et al.*, 2018). We use Receiver Operating Characteristic (ROC) curves to assess the performance of ranking differentially expressed genes using the log likelihood ratio (LLR) against a constant model. For the low dispersion data both the NB and Gaussian likelihood perform similarly well for low and high expression levels but the Poisson likelihood does not perform as well in the high expression case. For the high dispersion data the NB likelihood performs best and is clearly superior to both the Gaussian and Poisson likelihood. Anscombe’s transformation does not improve the performance of the Gaussian likelihood model and typically makes it worse.

**Figure 2:**
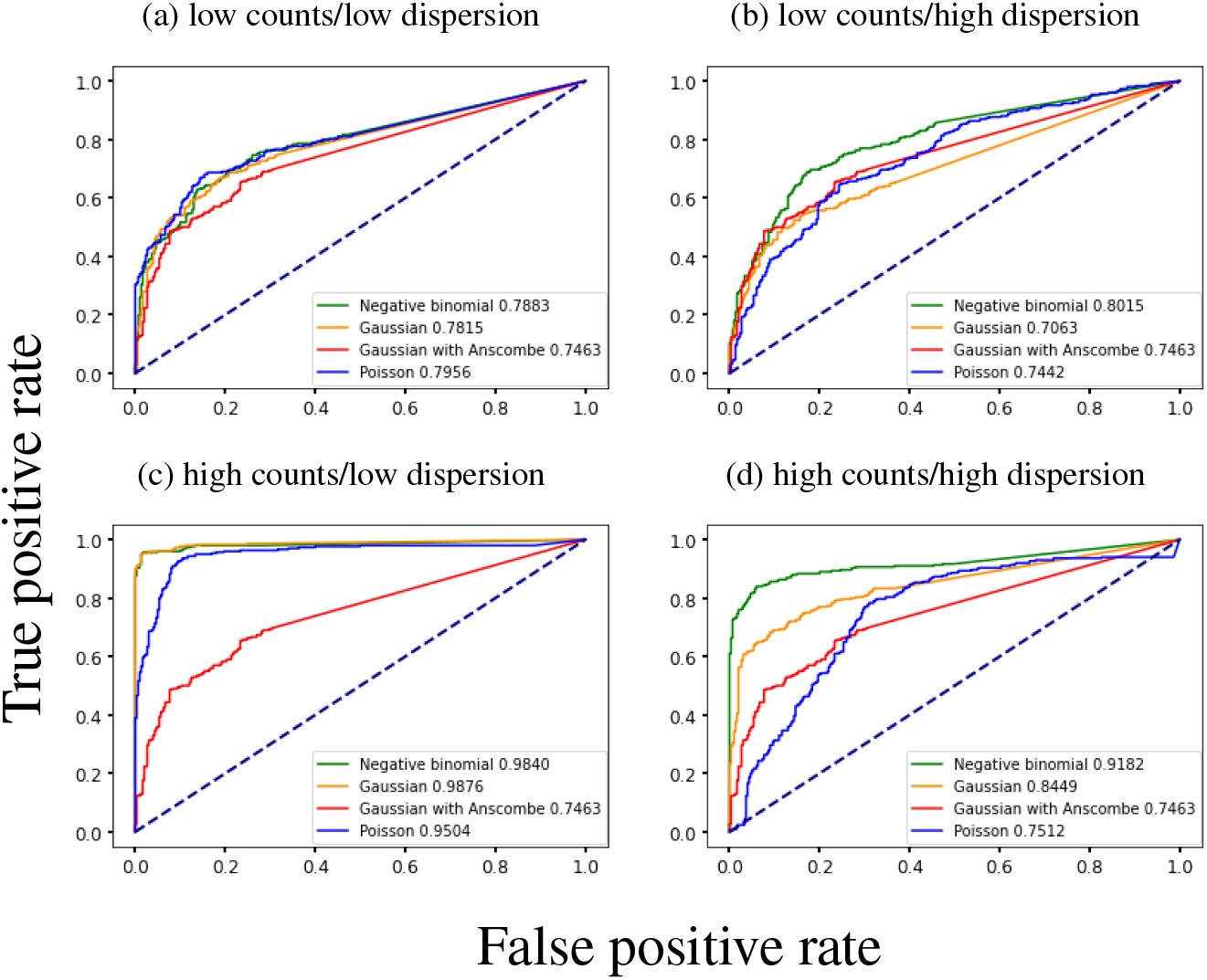
ROC curves for a one-sample test using a Gaussian process to identify dynamic time-series from synthetic counts data. Negative binomial, Gaussian and Poisson likelihoods are used to identify differentially expressed genes with a one-sample test. Genes are ranked by likelihood ratio between a dynamic and constant model. Results are shown for (a) low counts/low dispersion (b) low counts/high dispersion (c) high counts/low dispersion and (d) high counts/high dispersion datasets.

### 3.2 Assessment on tradeSeq cyclic time course data

The tradeSeq method uses spline-based generalized additive models (GAMs) with a NB likelihood to identify genes significantly associated with changes in pseudotime (Van den Berge *et al.*, 2020) and/or different lineages. The method was benchmarked against a number of other methods for identifying dynamic genes, including the GPfates method based on a GP regression, and was shown to perform better than other methods on benchmark data.

We compare the performance of GPcounts running a one-sample test with NB likelihood and Gaussian likelihood against tradeSeq using an association test on ten cyclic single-cell simulation datasets from the tradeSeq benchmark. The datasets were simulated using the dynverse toolbox (Saelens *et al.*, 2019) where the number of genes are between 312 – 444, number of cells are 505 – 507 and the percentages of differentially expressed genes are between 42 – 47%. The tradeSeq package uses a trajectory-based differential expression analysis method that uses a generalized additive model (GAM) with NB likelihood where each lineage is modeled as a cubic spline function. tradeSeq requires cell weight information to assign each cell to a lineage and packages such as slingshot (Street *et al.*, 2018) or GPfates (Lönnberg *et al.*, 2017) can be used to assign cell weight information. In Figure 3 (a) we compare the average performance of GPcounts and tradeSeq using inferred pseudotime trajectories on the ten simulated datasets from (Van den Berge *et al.*, 2020). In Figure 3 (b) we compare against ground truth time trajectories on the same data. The results in Figure 3 (a) suggest that tradeSeq performs better than GPcounts. However, when looking at specific examples we found that many constant examples that were incorrectly classified as dynamic by GPcounts appear to have dynamic profiles. Comparing these pseudotime profiles with profiles against the ground truth time shows that the dynamic structure is actually an artifact introduced by inferring pseudotime (in this case using slingshot). Figure 2 in the Supplementary Material shows an example of gene *H*1672 from dataset1 fitted using GPcounts with an NB likelihood where in (a) the pseudotime is estimated using slingshot and in (b) the true time is shown. Figure 3 (b) shows that the average performance of GPcounts with NB or Gaussian likelihoods is better than using tradeSeq when using the ground truth time information, and this remains the case for different numbers of knots [3, 5,10] (the number selected automatically by the package is 5). Detailed performance on each dataset using pseudotime information and using the true time information can be found in the Supplementary Material.

**Figure 3:**
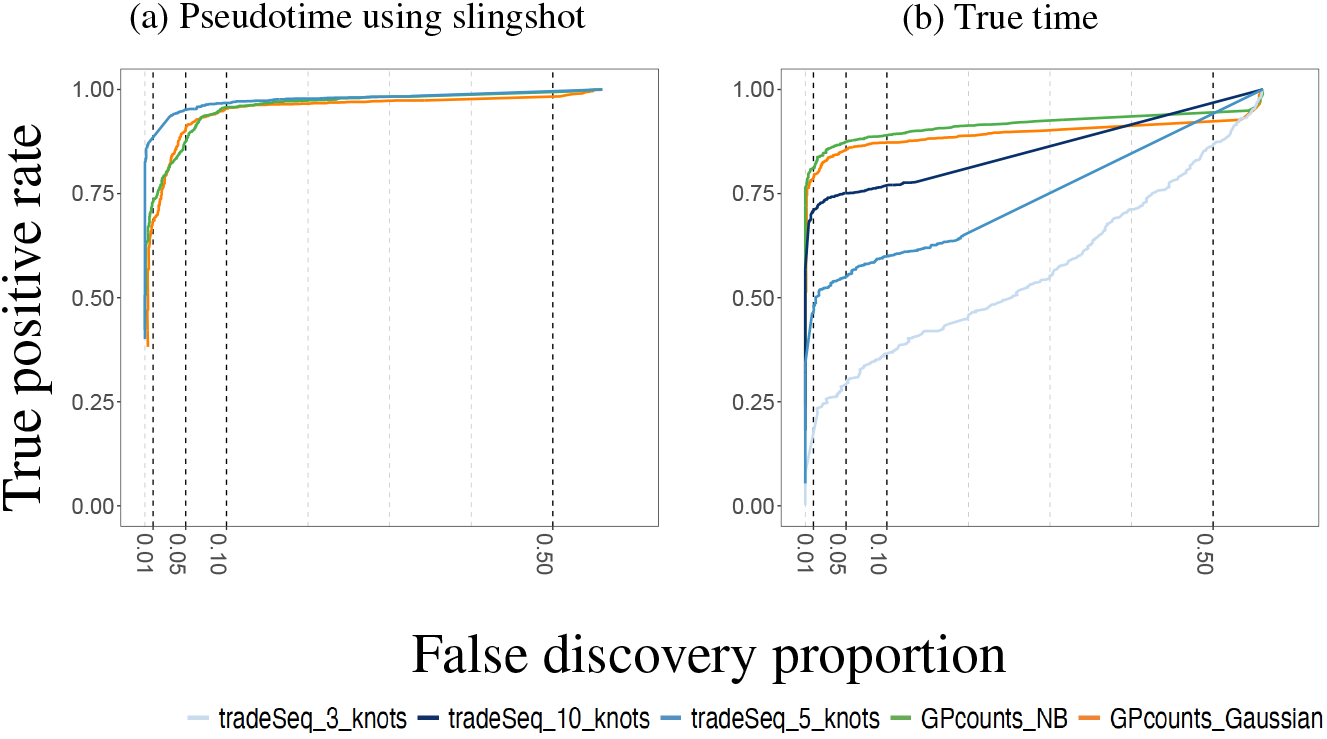
Mean performance of GPcounts running a one-sample test with NB likelihood and Gaussian likelihood versus the tradeSeq method running an association test over ten cyclic datasets from Van den Berge *et al.* (2020). In (a) we show results using pseudotime profiles with pseudotime estimated using slingshot. In (b) we show results using profiles against the true time used to generate the simulated data.

For scRNA-seq data the number of cells can be very large and computational efficiency becomes important. We benchmark the computation time for a one-sample test with GPcounts on the tenth synthetic cyclic dataset which has 312 genes and N = 507 cells. We use the full GP and compare it with a more computationally efficient sparse GP with different numbers of inducing points *M*, using the M-DPP algorithm (Burt *et al.*, 2020) to set the locations of inducing points. We choose different percentages of *N* to set the number of inducing points *M*. Our results in Figure 4 show that using sparse version gives results highly correlated with the full version dataset ranked by LLR while computation time reduces with *M*. Choosing the default *M* = 5%(*N*) = 25 inducing points decreases the computational time by ≈ 88% while obtaining a 96% Spearman correlation score between the LLRs.

**Figure 4:**
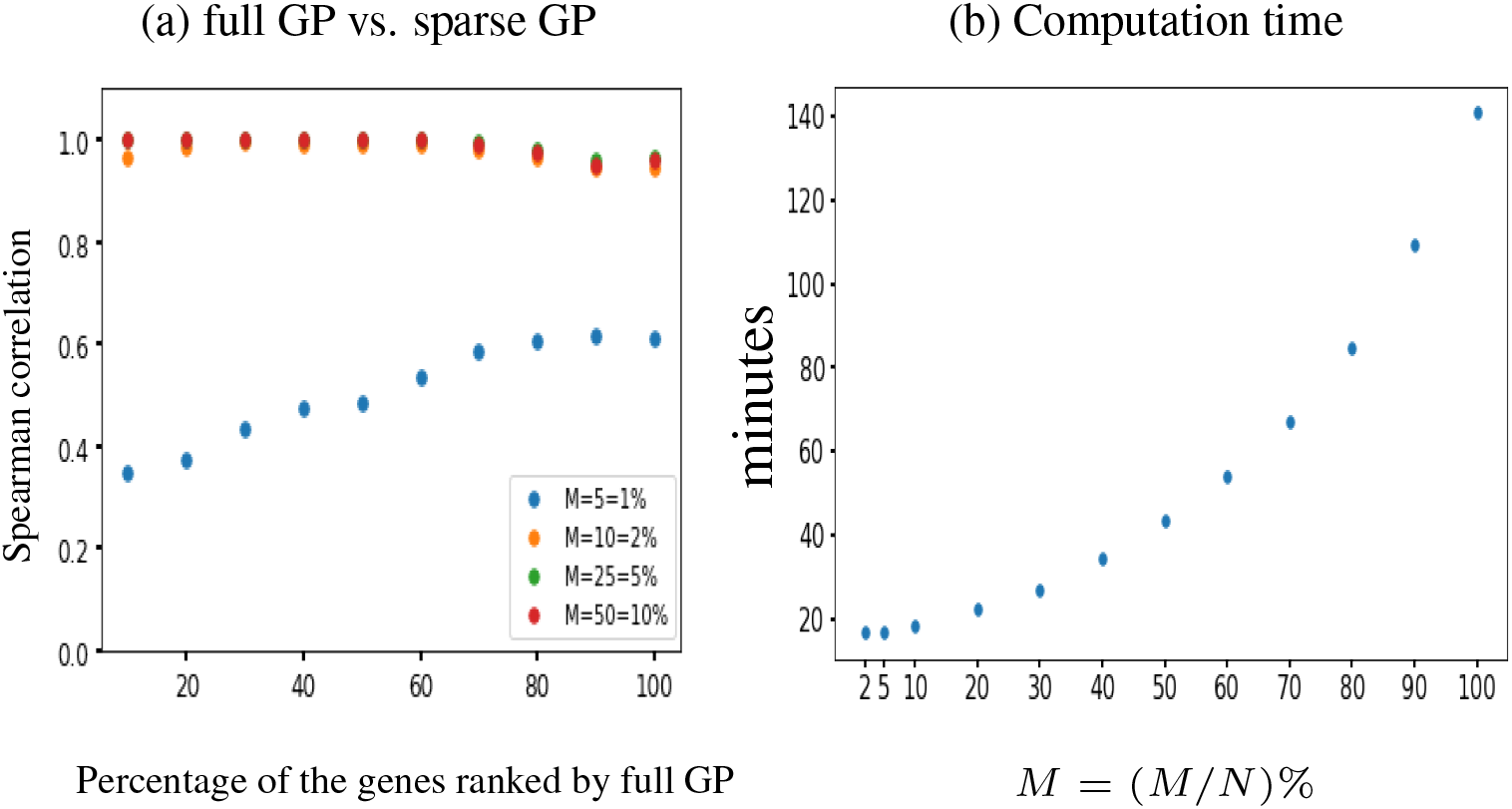
The performance of GPcounts with NB likelihood using full GP versus sparse version with different number of inducing points M on the tenth synthetic cyclic scRNA-seq dataset (Van den Berge *et al.*, 2020). Spearman correlation scores for the dataset ranked by full GP LLR score shown in (a) at different choices of M and the computation time in minutes shown in (b).

### 3.3 Modelling scRNA-seq pseudotime-series

We apply GPcounts on mouse pancreatic *α* cell data from scRNA-seq experiments without UMI normalisation (Qiu *et al.*, 2017) which has large numbers of zeros for some genes. We use the pseudotime inference results from Qiu *et al.* (2017) which are based on PCA. Figure 5 shows the inferred trajectory for two genes with many zero count measurements: Fam184b with 86% zero counts and Pde1a with 68% zero counts. From left to right we show the GP regression fit with Gaussian and NB likelihoods respectively. For both genes the Gaussian model is unable to effectively model the high probability region at zero counts, due to the symmetric nature of the distribution. In the Supplementary Material we run GPcounts with ZINB likelihood and compare it to NB and Gaussian likelihoods. Our results show that the ZINB model is a little better calibrated for Fam184b (Supplementary Figure 9). However, the fits for NB and ZINB are very similar for most genes (Supplementary Figure 10).

**Figure 5:**
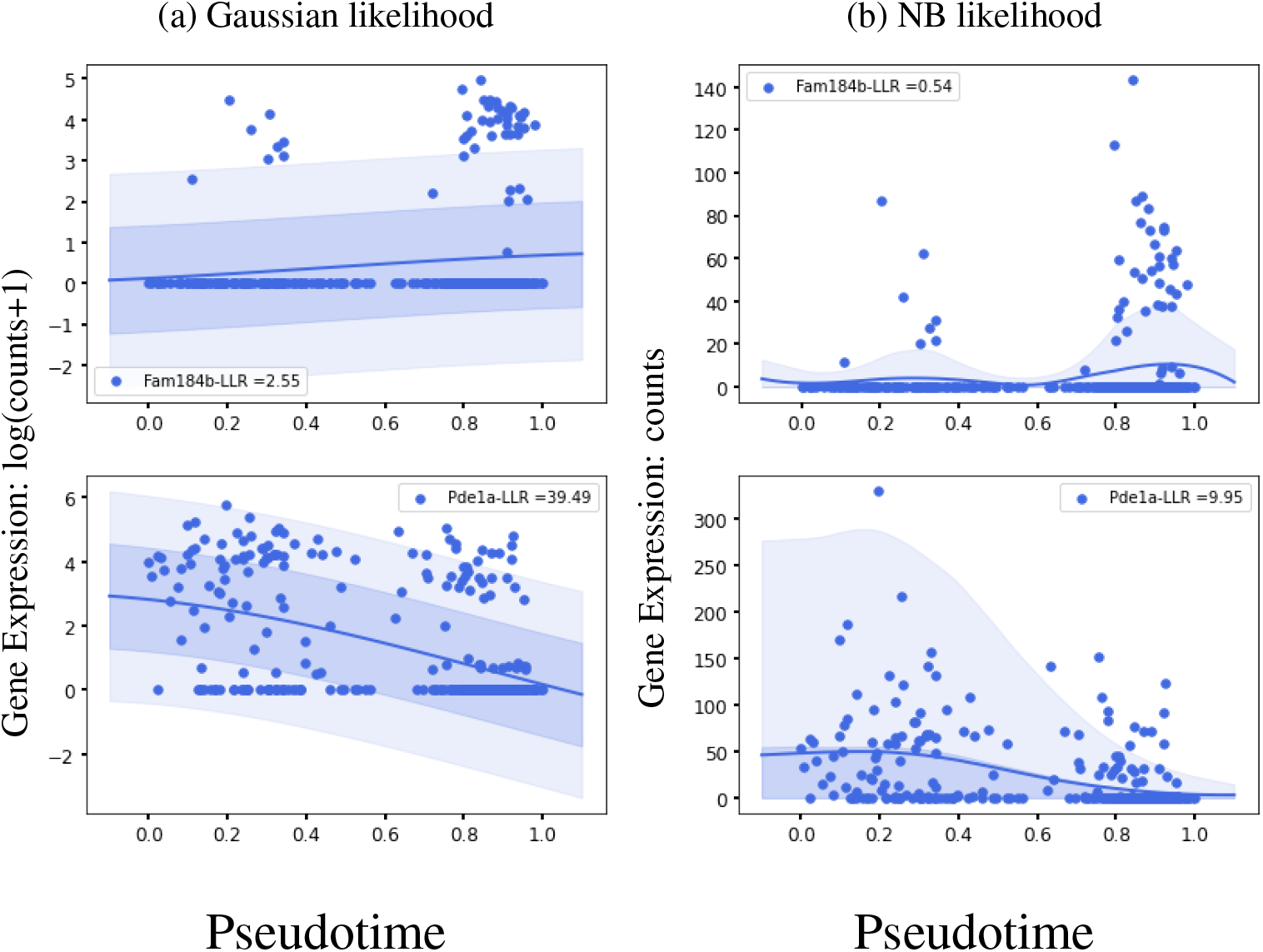
GP models of gene expression against pseudotime (Qiu *et al.*, 2017). We show two genes with a large number of zero-counts: Fam184b with 86% zeros in the first row and Pde1a with 68% zeros in the second row. We show the inferred mean trajectory and credible regions using (a) Gaussian likelihood and (b) NB likelihood.

Qiu *et al.* (2017) identify 611 differentially expressed genes in mouse pancreatic *α* cells using DESeq2 (Love *et al.*, 2014) which they applied to two distinctive clusters of cells along the pseudotime dimension. Since DESeq2 also assumes an NB model, we examined whether the results from GPcounts would be closer to DESeq2 than to a Gaussian GP model. We ran DESeq2 with time as a covariate and used the adjusted *p*-values as a score to label differentially expressed genes. We then compared the performance of one-sample tests using the NB likelihood and Gaussian likelihood. We consider different adjusted *p*-value thresholds to identify DE genes according to DESeq2 and look at the concordance with using a GP with NB and Gaussian likelihoods. Figure 6 shows precision–recall curves to explore how similarly GPcounts performs compared to DESeq2. We see that with the NB likelihood the genes obtained are very similar to those found by DESeq2. With a Gaussian likelihood the GP identifies very similar genes among its top-ranked DE genes but has much less concordance further down the list. This result is also reflected in the higher rank correlation between the DESeq and NB likelihood GP results than with the Gaussian likelihood GP (Figure 7). This suggests that the test statistic is more influenced by the form of likelihood term rather than the form of regression model in this example. However, for datasets with cells more evenly distributed across pseudotime the form of the regression model plays a more important role (e.g. as in Section 3.2).

**Figure 6:**
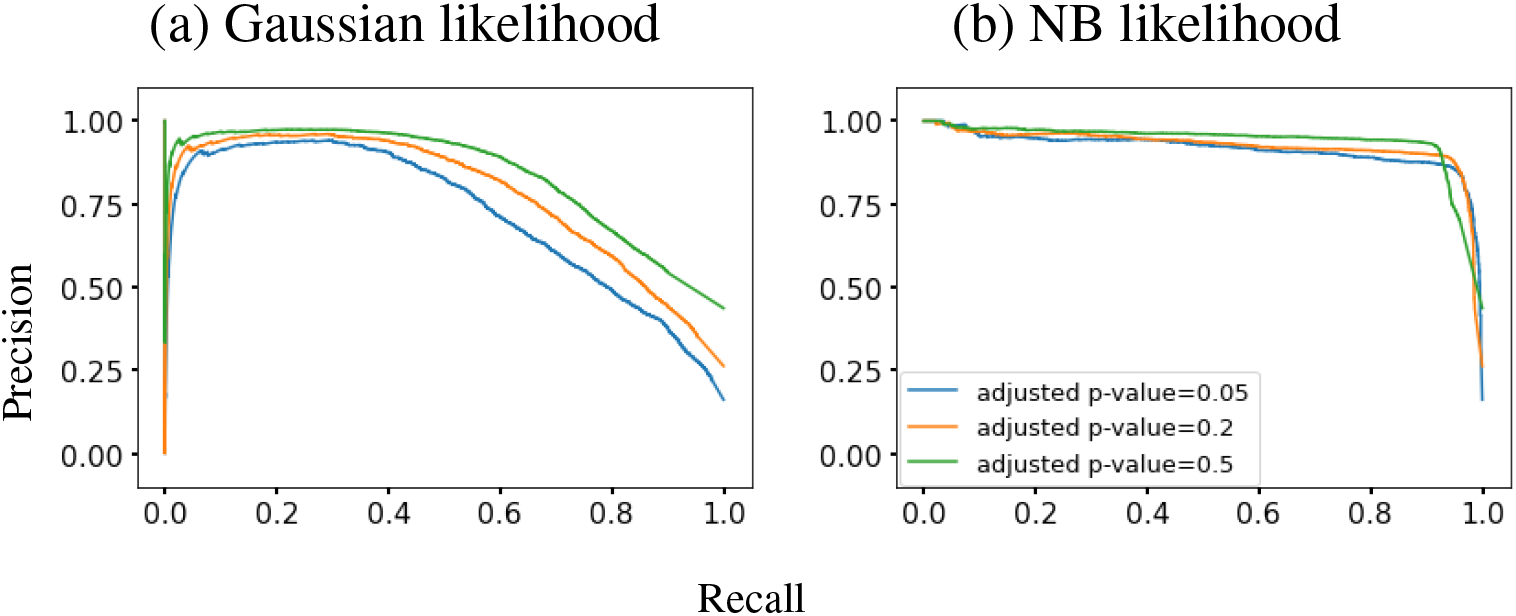
Precision-recall curves for GPcounts one-sample test with (a) Gaussian likelihood and (b) NB likelihood assuming ground truth from DESeq2 with different adjusted *p*-value thresholds. Mouse pancreatic *α*-cell scRNA-seq data from Qiu *et al.* (2017).

**Figure 7:**
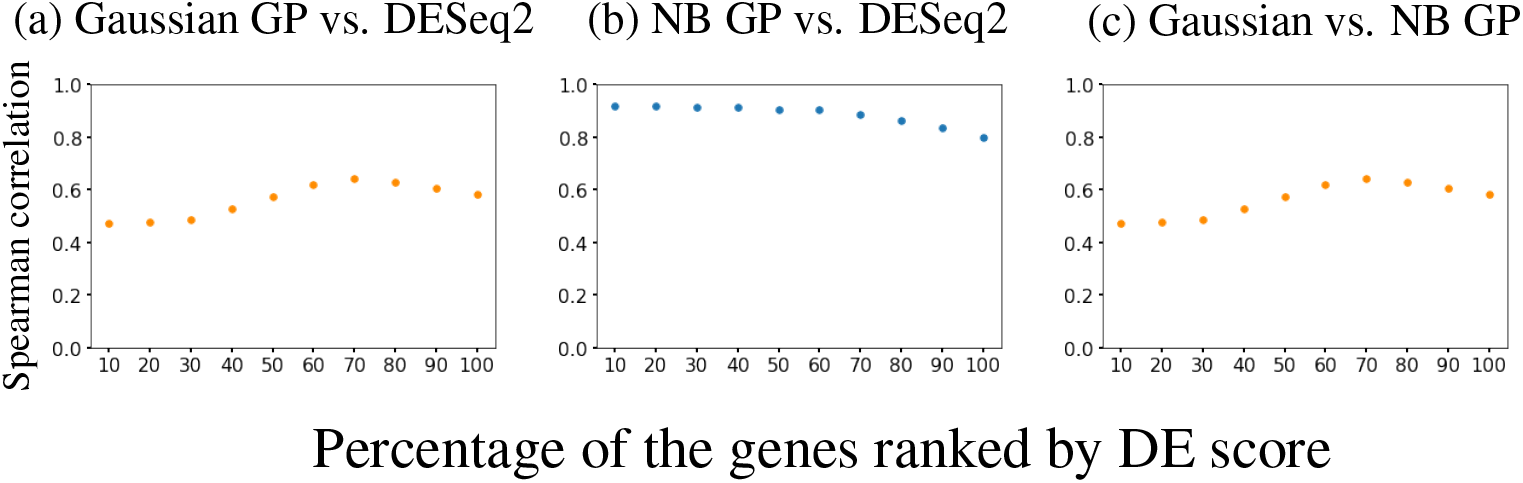
Spearman correlation scores for different percentages of the mouse pancreatic *α*-cell scRNA-seq data (Qiu *et al.*, 2017) ranked by DESeq2 adjusted *p*-value in (a) and (b) and by NB GP LLR in (c). We show (a) Gaussian likelihood GP versus DESeq2, (b) NB likelihood GP versus DESeq2 and (c) Gaussian likelihood GP versus NB likelihood GP.

In the Supplementary Material we show results using the sparse approximation for more efficient computational inference and show that the the M-DPP algorithm for setting the inducing point locations (Burt *et al.*, 2020) provides much better performance on this dataset than using k-means clustering.

### 3.4 Identification of spatially variable genes

We applied GPcounts to spatial transcriptomics sequencing counts data from mouse olfactory bulb (Ståhl *et al.*, 2016) consisting of measurements of 16 218 genes at 262 spatial locations. We compare performance to SpatialDE which uses a GP with a Gaussian likelihood (Svensson *et al.*, 2018) and we use as similar a set-up as possible in order to make the comparison fair. We filter out genes with less than three total counts and spatial locations with less than ten total counts and ended up with 14 859 genes across 260 spatial locations. To assess statistical significance we used the q-value method (Storey and Tibshirani, 2003) to determine an FDR cut-off (0.05) based on p-values assuming a χ^2^-distribution of LLRs. We compare our findings in Figure 8(a), where GPcounts identified 1096 spatially variable (SV) genes in total, whilst SpatialDE identified 67, with 63 of them overlapping between the two methods. It is worth pointing out here that, in the current SpatialDE version, the log likelihood statistic is incorrect, as the LLR rather than twice the LLR is used in the test. Correcting this error leads to 345 SV genes being called at 5% FDR threshold. However, GPcounts still identifies many more SV genes. We also compare with the SPARK method (Sun *et al.*, 2020) which is based on a GP with a Poisson likelihood that models over-dispersion using a white noise kernel. The SPARK and Trendsceek methods, which consider calibrated *p*-values under a permuted null, identified 772 and 0 SV genes respectively (Sun *et al.*, 2020). In Figure 8(a) we compare SPARK to GPcounts and SpatialDE and we find that GPcounts identifies more SV genes at the same FDR level.

**Figure 8:**
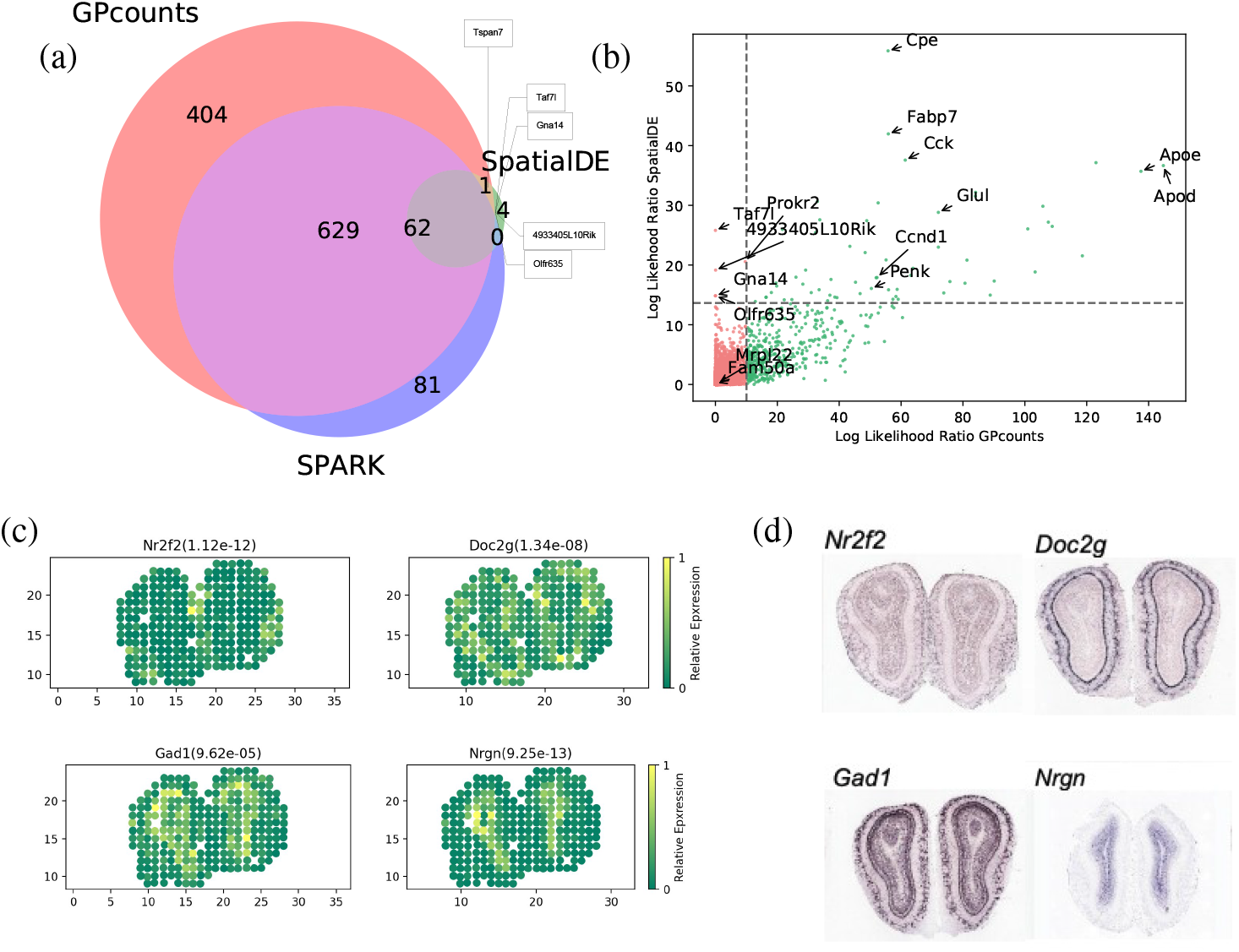
(a) Venn diagram shows the overlap between GPcounts’s, SpatialDE’s and SPARK’s spatially variable (SV) genes with the green area showing the four SV genes that are only identified by SpatialDE. (b) GPcounts’s LLR versus SpatialDE’s LLR. The dashed horizontal and vertical lines correspond to 5% FDR threshold. (c) Relative spatially resolved expression profiles for four selected SV genes, with their *q*-value in the parenthesis. (d) In-situ hybridization images of the selected SV genes in (c). The images are taken from Ståhl *et al.* (2016).

It is possible that the χ^2^-distribution is not perfectly calibrated and therefore we also implemented a permutation test approach where we randomly rearrange spatial coordinates to calculate *p*-values based on the permuted null. Under the permuted null, GPcounts indentified 1202 SV genes at the 5% FDR threshold, a much higher number than both the SpatialDE and SPARK methods (Supplementary Figure 13).

In Figure 8(b) we plot the GPcounts LLR versus SpatialDE LLR, showing in quadrants the genes that are identified as SV by each method. Genes Olfr635, Gna14, Taf7l and 4933405L10Rik were only identified by SpatialDE as significant. However, these genes were all extremely low expressed with three having the minimum number of counts (three) after filtering, while two of those are expressed in only one location (Supplementary Figure 13(c)). Relative expression profiles of four selected genes detected by GPcounts as SV are illustrated in Figure 8(c) with their *q*-values. Their spatial patterns match the associated profiles obtained with in-situ hybridization in the Allen Brain Atlas (Fig. 8(d)).

Checking against ten biologicaly important marker genes, known to be spatially expressed in the mitral cell layer (Ståhl *et al.*, 2016), GPcounts identified nine of those (Doc2g, Slc17a7, Reln, Cdhr1, Sv2b, Shisa3, Plcxd2, Nmb, Rcan2) while SpatialDE identified three (Doc2g, Cdhr1, Slc17a7) and SPARK identified eight (the same as GPcounts but missing Sv2b). Supplementary Figure 14(a) shows relative expression profiles of three selected marker genes (Fabp7, Rbfox3 and Eomes) and one house keeping gene (Actb) detected by GPcounts and SPARK as SV. None of these four genes is identified as SV by SpatialDE. Their associated profiles obtained with in-situ hybridization in the Allen Brain Atlas are shown in Supplementary Figure 14(b).

### 3.5 Identifying gene-specific branching locations

We used scRNA-seq of haematopoietic stem cells (HSCs) from mouse (Paul *et al.*, 2015) to demonstrate the identification of branching locations for individual genes. The data contain cells that are differentiated into myeloid and erythroid precursor cell types. Paul *et al.* (2015) analysed changes in gene expression for myeloid progenitors and created a reference compendium of marker genes related to the development of erythrocytes and several other types of leukocytes from myeloid progenitors. The Slingshot algorithm is used to get trajectory-specific pseudotimes as well as assignment of cells to different branches. After removing two small outlier cell clusters related to the dendritic and eosinophyl cell types, Slingshot infers two lineages for this dataset. These two outlier cell clusters do not belong to any particular lineage and have previously been excluded from trajectory inference and downstream analysis (Van den Berge *et al.*, 2020), which leaves us 2660 cells under consideration.

Figure 9 shows examples of GPcounts model fits with associated credible regions (upper sub-panels) as well as the posterior probability distribution over branching time (bottom sub-panels) for an early branching known bio-marker MPO (upper row) and for a late branching gene LY6E (bottom row). Here, we have sub-sampled the data and larger markers in Figure 9 represent the cells used in the inference process. Figure 9 (a) and (c) show the results with a Gaussian likelihood, equivalent to the model in Yang *et al.* (2016), while Figure 9 (b) and (d) show the results with the NB likelihood. In all cases, the bottom sub-panels reflect the significant amount of uncertainty associated with the identification of precise branching points. Both models provide a reasonably similar posterior probability of the branching time. However, looking at the credible region of the data we find that the model with NB likelihood better models the data. In the case of the Gaussian likelihood the credible regions are wide but they still miss some points that have zero values. In the case of NB likelihood the credible regions can adequately model the points having zero values. Further example fits are shown in the accompanying notebook.

**Figure 9:**
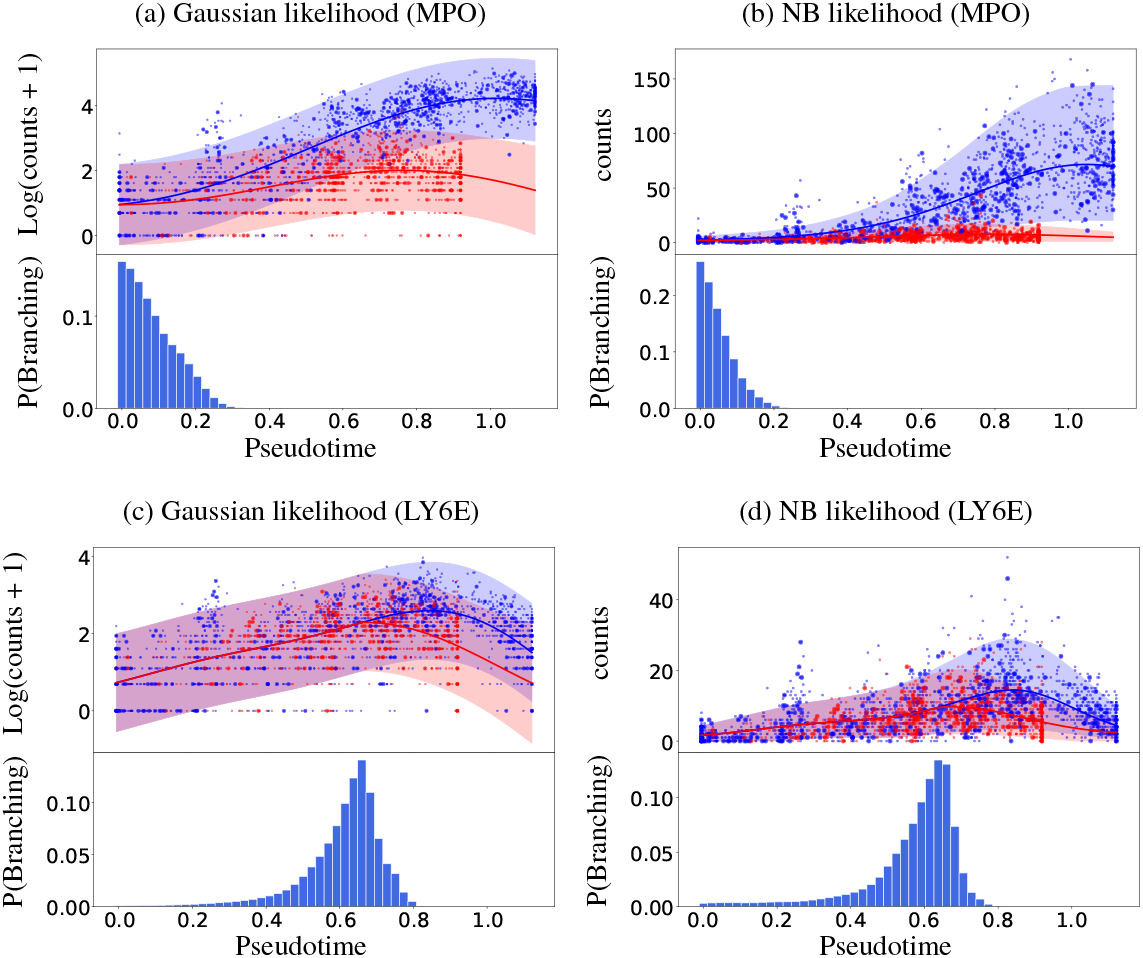
Mouse haematopoietic stem cells (Paul *et al.*, 2015): Identifying the gene-specific branching location for an early branching gene MPO (a and b) and a later branching gene LY6E (c and d). Cell assignments and pseudotime are estimated using the Slingshot algorithm. The upper sub-panels shows the GP regression fit based on the MAP estimate of the gene-specific differentiating time and the bottom sub-panels shows the posterior distribution over the differentiating or branching times. Bigger markers used in the upper sub-panels indicate the sub-sampled cells used in the inference. Results on the left are for the Gaussian likelihood fit (a,c) and on the right for the NB likelihood fit (b,d).

It should be noted that trajectory-based inference can give different results depending on the non-linear relationship between pseudotime and real time, which can differ along each lineage. Time warping methods can therefore be useful to obtain improved identification of branching trajectories (Alpert *et al.*, 2018).

## 4 Conclusion

We have developed a Gaussian process (GP) regression method, GPcounts, implementing a negative binomial (NB) likelihood in GP inference. This provides a useful tool for RNA-seq data from time-series, single-cell and spatial transcriptomics experiments. Our results show that the NB likelihood can provide a substantial improvement over a Gaussian likelihood when modelling counts data. Our simulations suggest that gains are largest when data are highly over-dispersed. For lower dispersion data the performance of the Gaussian and NB likelihood is similar. We find that the Poisson distribution likelihood performs very poorly for highly expressed genes even for relatively low dispersion.

In terms of our example applications the most notable success was in the application to spatial transcriptomics data. We found a substantial difference using the NB likelihood compared to the SpatialDE method that is based on a Gaussian likelihood GP. Using a similar normalisation and testing set-up, we found a much larger set of spatially variable (SV) genes than SpatialDE. Similarly, we found more SV genes than the over-dispersed Poisson method, SPARK, which also uses GP inference but with differences in both the modelling and inference set-up. When modelling the scRNA-Seq data from Qiu *et al.* (2017) against pseudotime we found that the NB GP identifies DE gene lists more similar to DESeq2 than to a Gaussian GP. This suggests that the likelihood function plays a more important role than the regression model in some problems, emphasising the importance of using an appropriate likelihood function. Finally, we applied a branching kernel to infer the initial point when the two gene expression time profiles begin to diverge along a pseudotime trajectory. For genes with strong branching evidence the NB and Gaussian likelihood provided similar inference results in this case.

To improve the practical performance of GPcounts we implement a heuristic to detect locally optimal solutions and to detect numerical instability. Since the naive GP scales cubically with number of time points we improve the computational requirements through a sparse inference algorithm from the GPflow library (De G. Matthews *et al.*, 2017) using the M-DPP algorithm for learning the inducing points (Burt *et al.*, 2020). Our implementation of GPcounts is flexible and can easily be extended to work with any kernel or likelihood compatible with the GPflow library. A natural next step would be to better parellelise model fitting for each gene.

## References

Ahmed, S., Rattray, M., and Boukouvalas, A. (2019). GrandPrix: scaling up the Bayesian GPLVM for single-cell data. Bioinformatics, 35(1), 47–54.

Äijö, T., Butty, V., Chen, Z., Salo, V., Tripathi, S., Burge, C. B., Lahesmaa, R., and Lähdesmäki, H. (2014). Methods for time series analysis of RNA-seq data with application to human Th17 cell differentiation. Bioinformatics, 30(12), i113–i120.

Alpert, A., Moore, L. S., Dubovik, T., and Shen-Orr, S. S. (2018). Alignment of single-cell trajectories to compare cellular expression dynamics. Nature methods, 15(4), 267.

Anscombe, F. J. (1948). The transformation of Poisson, binomial and negative-binomial data. Biometrika, 35(3/4), 246–254.

Arnol, D., Schapiro, D., Bodenmiller, B., Saez-Rodriguez, J., and Stegle, O. (2019). Modeling Cell-Cell Interactions from Spatial Molecular Data with Spatial Variance Component Analysis. Cell Reports, 29(1), 202–211.e6.

Bauer, M., van der Wilk, M., and Rasmussen, C. E. (2016). Understanding probabilistic sparse Gaussian process approximations. In Advances in neural information processing systems, pages 1533–1541.

Bernardo, J., Bayarri, M., Berger, J., Dawid, A., Heckerman, D., Smith, A., West, M., et al. (2003). The variational bayesian em algorithm for incomplete data: with application to scoring graphical model structures. Bayesian statistics, 7(453-464), 210.

Boukouvalas, A., Hensman, J., and Rattray, M. (2018). BGP: identifying gene-specific branching dynamics from single-cell data with a branching Gaussian process. Genome biology, 19(1), 65.

Burt, D. R., Rasmussen, C. E., and van der Wilk, M. (2020). Convergence of sparse variational inference in gaussian processes regression. Journal of Machine Learning Research, 21(131), 1–63.

Choi, K., Chen, Y., Skelly, D. A., and Churchill, G. A. (2020). Bayesian model selection reveals biological origins of zero inflation in single-cell transcriptomics. bioRxiv.

Cleveland, W. S. (1979). Robust locally weighted regression and smoothing scatterplots. Journal of the American statistical association, 74(368), 829–836.

De G. Matthews, A. G., Van Der Wilk, M., Nickson, T., Fujii, K., Boukouvalas, A., León-Villagrá, P., Ghahramani, Z., and Hensman, J. (2017). GPflow: A Gaussian process library using TensorFlow. The Journal of Machine Learning Research, 18(1), 1299–1304.

Edsgärd, D., Johnsson, P., and Sandberg, R. (2018). Identification of spatial expression trends in single-cell gene expression data. Nature Methods, 15(5), 339–342.

Hensman, J., Lawrence, N. D., and Rattray, M. (2013). Hierarchical bayesian modelling of gene expression time series across irregularly sampled replicates and clusters. BMC bioinformatics, 14(1), 252.

Hensman, J., Matthews, A., and Ghahramani, Z. (2015). Scalable variational gaussian process classification. In Artificial Intelligence and Statistics, pages 351–360. PMLR.

Kalaitzis, A. A. and Lawrence, N. D. (2011). A simple approach to ranking differentially expressed gene expression time courses through Gaussian process regression. BMC bioinformatics, 12(1), 180.

Leong, H. S., Dawson, K., Wirth, C., Li, Y., Connolly, Y., Smith, D. L., Wilkinson, C. R., and Miller, C. J. (2014). A global non-coding rna system modulates fission yeast protein levels in response to stress. Nature communications, 5(1), 1–10.

Lönnberg, T., Svensson, V., James, K. R., Fernandez-Ruiz, D., Sebina, I., Montandon, R., Soon, M. S., Fogg, L. G., Nair, A. S., Liligeto, U., et al. (2017). Single-cell rna-seq and computational analysis using temporal mixture modelling resolves th1/tfh fate bifurcation in malaria. Science immunology, 2(9).

Love, M. I., Huber, W., and Anders, S. (2014). Moderated estimation of fold change and dispersion for RNA-seq data with DESeq2. Genome biology, 15(12), 550.

McDowell, I. C.,Manandhar, D., Vockley, C. M., Schmid, A. K., Reddy, T. E., and Engelhardt, B. E. (2018). Clustering gene expression time series data using an infinite gaussian process mixture model. PLoS computational biology, 14(1), e1005896.

Millman, K. J. and Aivazis, M. (2011). Python for scientists and engineers. Computing in Science & Engineering, 13(2), 9–12.

Opper, M. and Archambeau, C. (2009). The variational gaussian approximation revisited. Neural computation, 21(3), 786–792.

Paul, F., Arkin, Y., Giladi, A., Jaitin, D. A., Kenigsberg, E., Keren-Shaul, H., Winter, D., Lara-Astiaso, D., Gury, M., Weiner, A., et al. (2015). Transcriptional heterogeneity and lineage commitment in myeloid progenitors. Cell, 163(7), 1663–1677.

Pierson, E. and Yau, C. (2015). ZIFA: Dimensionality reduction for zero-inflated single-cell gene expression analysis. Genome biology, 16(1), 241.

Qiu, W.-L., Zhang, Y.-W., Feng, Y., Li, L.-C., Yang, L., and Xu, C.-R. (2017). Deciphering pancreatic islet *β* cell and *α* cell maturation pathways and characteristic features at the single-cell level. Cell metabolism, 25(5), 1194–1205.

Rasmussen, C. E. and Williams, C. K. (2006). Gaussian process for machine learning. MIT press.

Risso, D., Perraudeau, F., Gribkova, S., Dudoit, S., and Vert, J.-P. (2018). A general and flexible method for signal extraction from single-cell rna-seq data. Nature communications, 9(1), 1–17.

Robinson, M. D. and Smyth, G. K. (2007). Moderated statistical tests for assessing differences in tag abundance. Bioinformatics, 23(21), 2881–2887.

Saelens, W., Cannoodt, R., Todorov, H., and Saeys, Y. (2019). A comparison of single-cell trajectory inference methods. Nature biotechnology, 37(5), 547–554.

Savitzky, A. and Golay, M. J. (1964). Smoothing and differentiation of data by simplified least squares procedures. Analytical chemistry, 36(8), 1627–1639.

Seeger, M. (2000). Bayesian model selection for support vector machines, gaussian processes and other kernel classifiers. In Advances in neural information processing systems, pages 603–609.

Ståhl, P. L., Salmén, F., Vickovic, S., Lundmark, A., Navarro, J. F., Magnusson, J., Giacomello, S., Asp, M., Westholm, J. O., Huss, M., Mollbrink, A., Linnarsson, S., Codeluppi, S., Borg, Å., Pontén, F., Costea, P. I., Sahlén, P., Mulder, J., Bergmann, O., Lundeberg, J., and Frisén, J. (2016). Visualization and analysis of gene expression in tissue sections by spatial transcriptomics. Science, 353(6294), 78–82.

Stegle, O., Denby, K. J., Cooke, E. J., Wild, D. L., Ghahramani, Z., and Borgwardt, K. M. (2010). Arobust Bayesiantwo-sampletestfor detecting intervals of differential gene expression in microarray time series. Journal of Computational Biology, 17(3), 355–367.

Storey, J. D. and Tibshirani, R. (2003). Statistical significance for genomewide studies. Proc Natl Acad Sci USA, 100(16), 9440–9445.

Street, K., Risso, D., Fletcher, R. B., Das, D., Ngai, J., Yosef, N., Purdom, E., and Dudoit, S. (2018). Slingshot: cell lineage and pseudotime inference for single-cell transcriptomics. BMC genomics, 19(1), 477.

Sun, S., Zhu, J., and Zhou, X. (2020). Statistical analysis of spatial expression patterns for spatially resolved transcriptomic studies. Nature Methods, 17(2), 193–200.

Svensson, V. (2020). Droplet scRNA-seq is not zero-inflated. Nature Biotechnology, pages 1–4.

Svensson, V., Teichmann, S. A., and Stegle, O. (2018). Spatialde: identification of spatially variable genes. Nature Methods, 15(5), 343–346.

Townes, F. W., Hicks, S. C., Aryee, M. J., and Irizarry, R. A. (2019). Feature selection and dimension reduction for single-cell RNA-Seq based on a multinomial model. Genome biology, 20(1), 1–16.

Van den Berge, K., De Bezieux, H. R., Street, K., Saelens, W., Cannoodt, R., Saeys, Y., Dudoit, S., and Clement, L. (2020). Trajectory-based differential expression analysis for single-cell sequencing data. Nature communications, 11(1), 1–13.

Yang, J., Penfold, C. A., Grant, M. R., and Rattray, M. (2016). Inferring the perturbation time from biological time course data. Bioinformatics, 32(19), 2956–2964.

