## Supplementary Material for "Non-parametric modelling of temporal and spatial counts data from RNA-seq experiments"

### Contents

**1 Zero-inflated negative binomial likelihood** **1**

**2 Practical considerations** **2**

**3 Assessment on tradeSeq cyclic time course data** **2**

**4 Identifying gene-specific branching locations** **2**

**5 Modelling scRNA-seq pseudotime-series** **8**

**6 Identification of spatially variable genes** **12**

#### 1 Zero-inflated negative binomial likelihood

In some cases the model may benefit from an excess of zeros beyond the negative binomial (NB) distribution assumption. For example, the NB distribution may not adequately model non-UMI scRNA-seq counts data since an excess of zeros may lead to an overestimate of the dispersion parameter and a biased mean estimate. Even in the case of UMI scRNA-seq data, the model may benefit from some additional flexibility when fitting data to an inferred pseudotime trajectory.

A zero-inflated distribution can be used to model count data with excessive zeros (Pierson and Yau, 2015; Risso *et al.*, 2018). These extra zeros are assumed to be generated by a separate process different from the negative binomial process that generates counts including zeros. The ZINB distribution is parametrised by $\Psi$ (the excess proportion of zeros) and the NB model parameters:

$$\mathrm{ZINB}(y;\mu,\alpha,\Psi) = \begin{cases} \Psi + (1-\Psi)\mathrm{NB}(y=0) \quad \text{for} \quad y=0\ , \\ (1-\Psi)\mathrm{NB}(y) \quad \text{for} \quad y \geq 1\ . \end{cases} \tag{1}$$

We parametrise the probability of dropout $\Psi$ using a Michaelis–Menten (MM) equation relating the probability of dropout to the mean of the NB distribution used to model the non-dropout data $\mu = e^{f(x)}$ (using the same link function as above for the NB likelihood),

$$\Psi = 1 - \frac{e^{f(x)}}{K_M + e^{f(x)}} \tag{2}$$

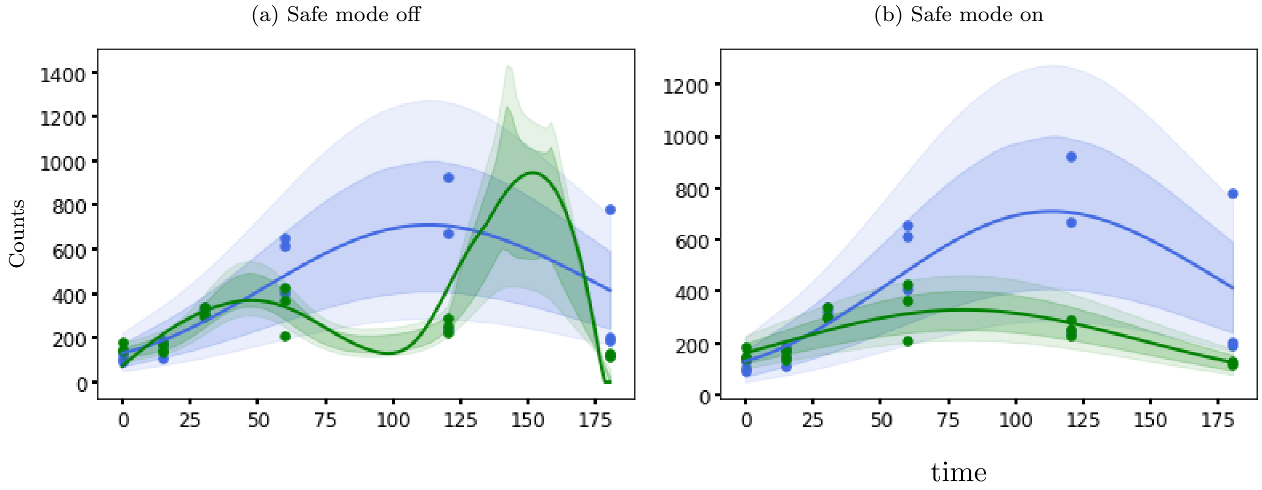


Figure 1: Gene SPAC11D3.01c from a two-sample time course RNA-seq experiment (Leong *et al.*, 2014). The GP is stuck in local optima in (a) with safe mode option switched off and GPcounts solved local optima with a random restart in (b) with safe mode option switched on.

where the Michaelis constant $K_M$ is estimated by our model. The MM equation used to model an enzyme reaction so it used to model the excess zeros which resulted from a failure in reverse transcription enzyme reaction (Andrews and Hemberg, 2016). Andrews and Hemberg (2016) shows that using MM equation to model excess zeros is better than using double exponential function used in ZIFA method (Pierson and Yau, 2015) and better than or similar to logistic regression.

#### 2 Practical considerations

We show an example of gene where the GP is stuck in local optima in figure 1 (a) fitted using GPcounts with NB likelihood and safe mode option is switched off while in (b) we show the same gene where the local optima is solved by random multiple restart when the safe mode option is switched on.

#### 3 Assessment on tradeSeq cyclic time course data

An example of non-differentially expressed gene $H1672$ from dataset1 fitted using GPcounts with NB likelihood. The pseudotime estimated using slingshot package in figure 2 (a) and we can see that using slingshot introduced temporal correlation between time points which led to the gene identified as differentially expressed gene by GPcounts. Using the true time information in figure 2 (b) shows no correlation between the time points and makes the gene to be identified by GPcounts as non-differentially expressed gene (see notebook accompanying the package for more examples).

The detailed performance of GPcounts with NB likelihood and Gaussian likelihood running one-sample test versus tradeSeq (Van den Berge *et al.*, 2020) running association test on each simulated dataset are shown in figure 3 and figure 4. Assuming pseudotime estimated using slingshot in figure 3 and using the true time information in figure 4.

#### 4 Identifying gene-specific branching locations

Paul *et al.* (2015) generated a list of biomarker genes for developing myeloid cells in their extensive analysis. The top six genes from their list are PRTN3, MPO, CAR2, CTSG, ELANE, and CAR1. They labelled all of these six genes as the “key genes” of hematopoiesis process. Thus these six genes are significantly differentially expressed (DE) between lineages. In fact it was found that MPO and CAR2 differentiated between erythroid progenitors and myeloid progenitors, where PRTN3 was identified as monocyte-specific, and the cluster of three genes ELANE, PRTN3 and MPO was found

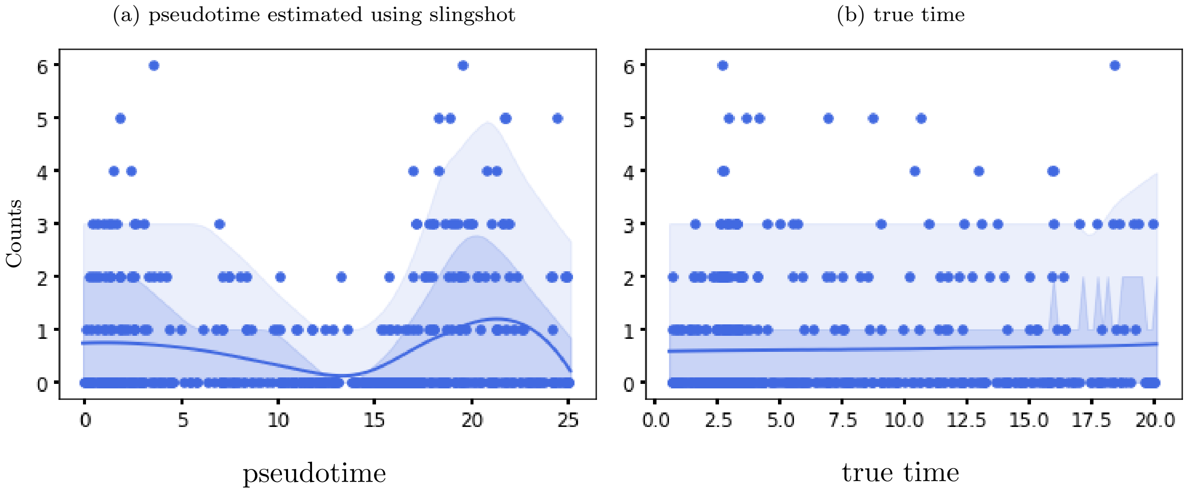


Figure 2: Gene $H1672$ is non-differentially expressed genes from dataset1 show slingshot introduced a temporal correlation between time points in (a) while the true time information show no correlation between time points in (b).

as the strongest markers for myeloid progenitors as well as monocytes (Van den Berge *et al.*, 2020). We have used the GPcounts model to investigate the branching dynamics of these six marker genes. Figure 5 and 6 show the examples of GP regression with NB likelihood (GPcounts model) and Gaussian likelihood models applied on the six biomarkers respectively. In all cases, the upper sub-panel shows the GP regression fit based on the MAP estimate of the gene-specific differentiating time and the lower sub-panel shows the posterior distribution over the differentiating or branching time point. The bigger markers used in the upper sub-panels to represent the sub-sampled cells that have been used in the inference. From the lower sub-panels, it is evident that both of these approaches can adequately accommodate the identification of genes having different expression patterns between lineages. However, the predictive distribution of the data depicted in upper sub-panels reflects that the GPcounts model, i.e. GP regression with NB likelihood better models the data. Single-cell data contain a lot of zeros. The credible regions are wide in the case of Gaussian likelihood and still miss some points that have zero values while the credible regions of the model with NB likelihood, i.e. the GPcounts model can adequately model the points having zero values.

Next, we investigate the robustness of this approach in identifying branching locations for the genes that are differentially expressed across the lineages, but show very little or no evidence of having different expression patterns at the end. IRF8, APOE and ERP29 are among the top genes having this type of expression patterns (Van den Berge *et al.*, 2020). These genes are known regulators of hematopoiesis and are discussed in previous studies (Shin *et al.*, 2011; Kurotaki *et al.*, 2014; Paul *et al.*, 2015; Murphy *et al.*, 2011). Identifying branching points for these genes is challenging as the model may be deceived easily, which may lead to wrong inference of branching locations.

Figure 7 (a) and (b) show the GPcounts model fit on the expression profile of genes IRF8 and APOE respectively. In each case, the upper sub-panel shows the GP regression model fit where the most probable branching location is identified using the MAP estimate; and the lower sub-panel describes the posterior probability over branching location. From Figure 7 it is evident that the proposed GPcounts model can identify gene-specific branching dynamics even when the gene expression patterns are very similar.

Figure 8 shows another example of the GPcounts model fit on the gene ERP29. The posterior probability of the branching time for gene ERP29 (lower sub-panel) indicates an interesting feature. The flat posterior distribution at the end of time range suggests there is not strong evidence that the gene is branching. To investigate that we have calculated the logged Bayes factor $r_g$. The posterior probability returned by the model (see Main paper) can be used to calculate the logged Bayes factor that provides evidence for whether the two time courses diverge (after any time) or are statistically indistinguishable. If the estimated branching time is closer to the start of the time course, it is more

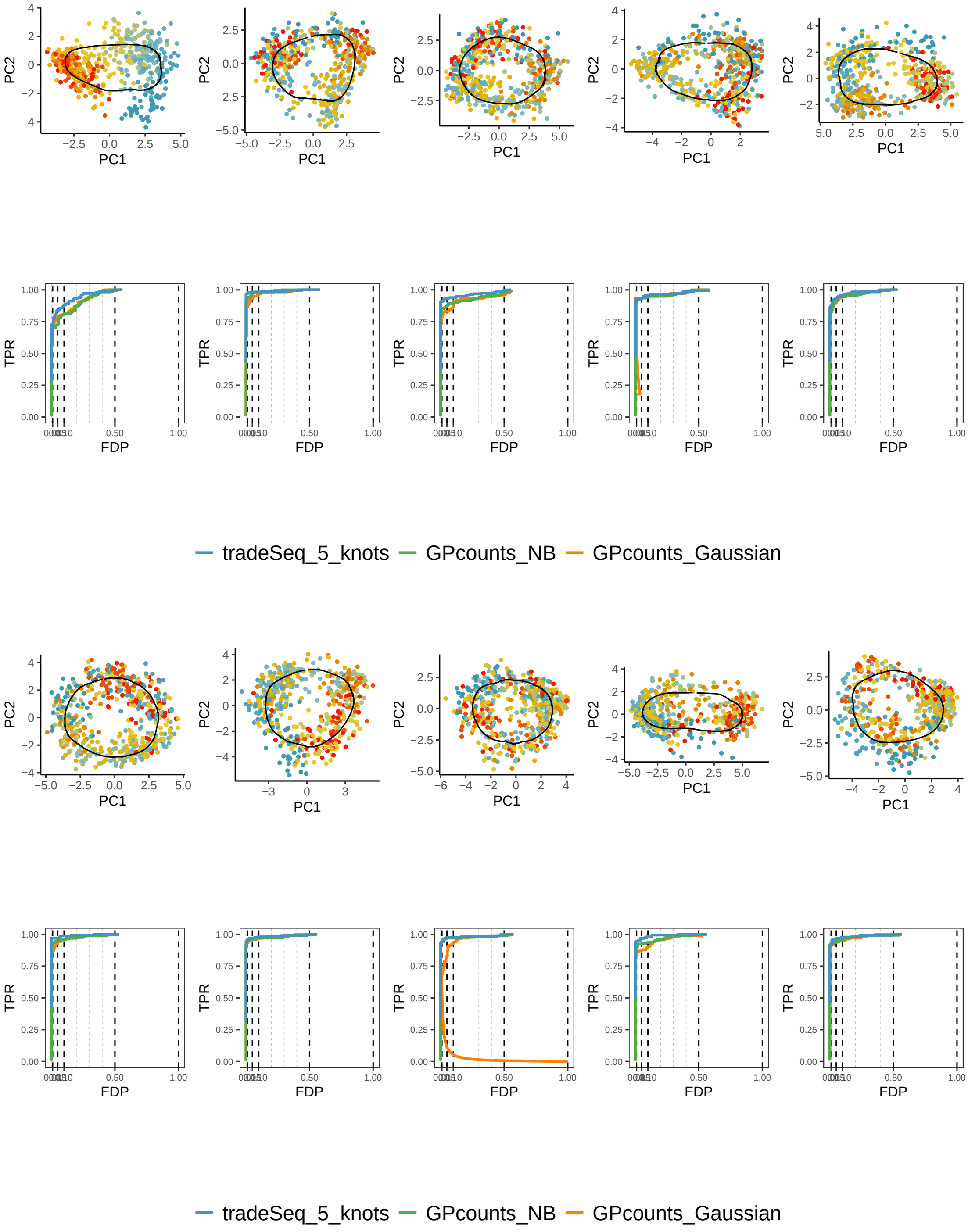


Figure 3: Ten simulated cyclic datasets. The first and the third rows show PCA plots where the second and fourth rows show the performance of GPcounts with NB likelihood and Gaussian likelihood versus tradeSeq using pseudotime estimated with slingshot.

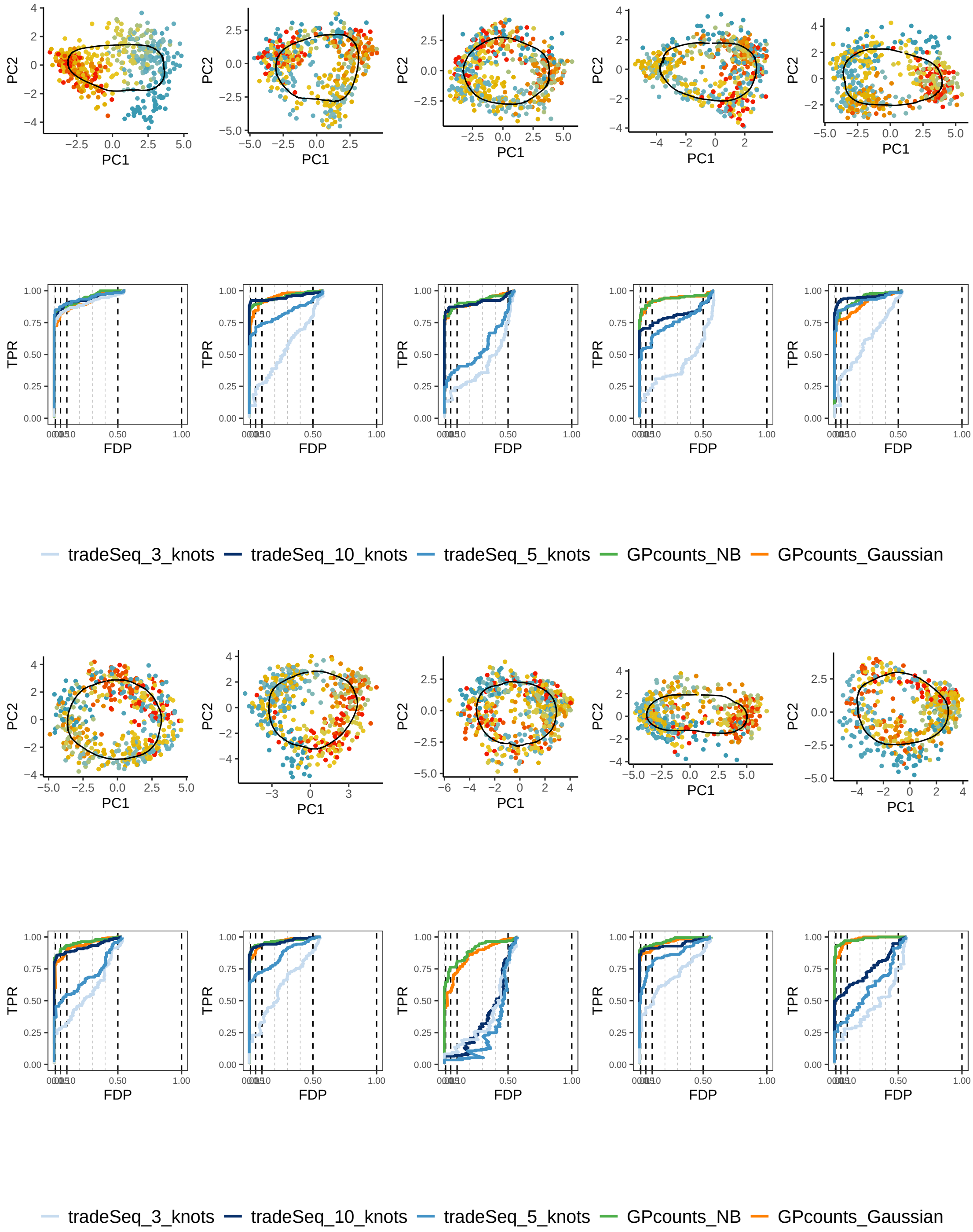


Figure 4: Ten simulated cyclic datasets. The first and the third rows show PCA plots where the second and fourth rows show the performance of GPcounts with NB likelihood and Gaussian likelihood versus tradeSeq using true time.

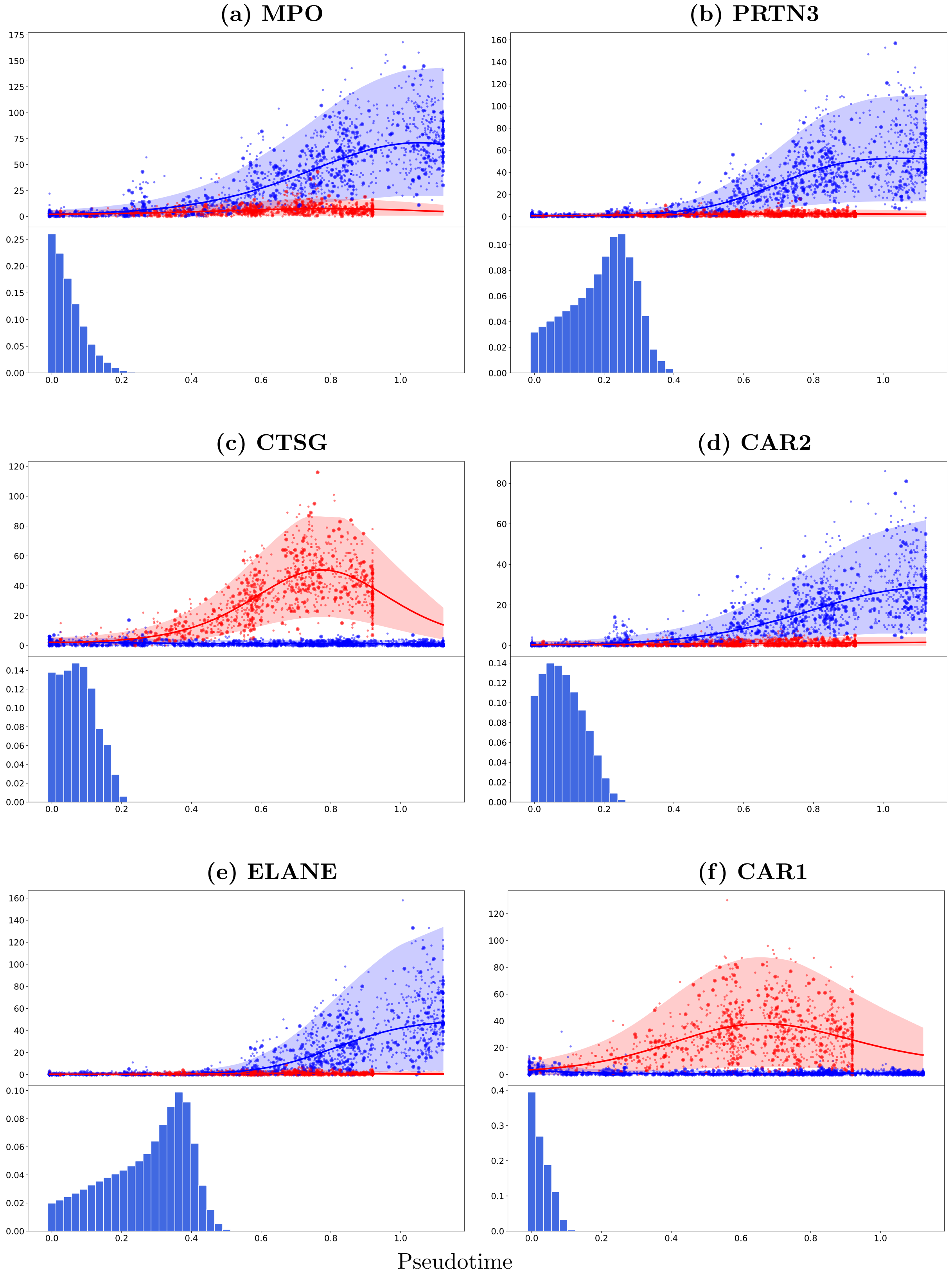


Figure 5: Mouse haematopoietic stem cells (Paul *et al.*, 2015): Examples of the GPcounts model (GP regression with Negative Binomial likelihood) fit on the top six biomarker genes for developing myeloid cells (upper sub-panels) and the posterior distribution over the branching times (lower sub-panels). Cell assignment to different branches as well as the pseudotime of each cell are calculated using the Slingshot algorithm. The bigger markers shown in upper sub-panels represent the sub-sampled cells that have been used in the inference. The GP regression fit depicted is based on the MAP estimate of the gene-specific differentiating or branching time. Lowess has been used to smooth the percentiles through pseudotime.

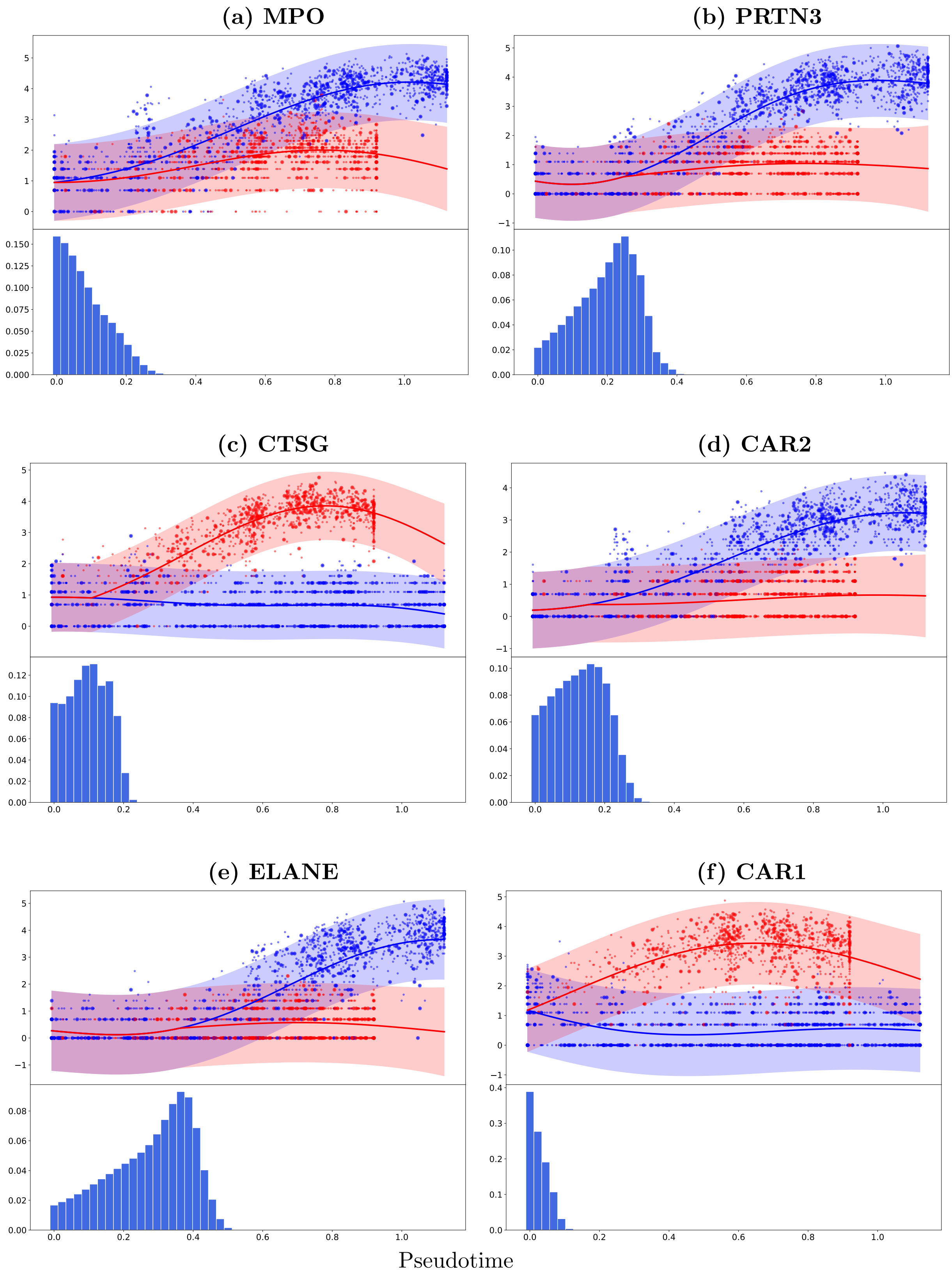


Figure 6: Mouse haematopoietic stem cells (Paul *et al.*, 2015): Examples of the GP regression model fit on the top six biomarker genes for developing myeloid cells (upper sub-panels) and the posterior distribution over the branching times (lower sub-panels). Cell assignment to different branches as well as the pseudotime of each cell are calculated using the Slingshot algorithm. The bigger markers shown in the upper sub-panels represent the sub-sampled cells that have been used in the inference. The GP regression fit depicted is based on the MAP estimate of the gene-specific differentiating or branching time.

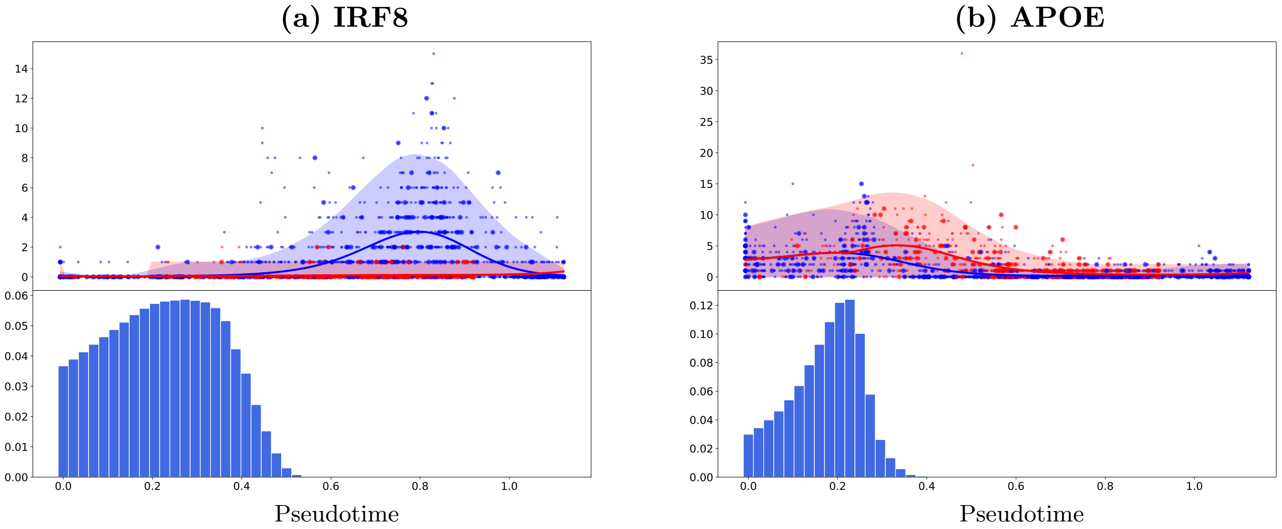


Figure 7: Mouse haematopoietic stem cells (Paul *et al.*, 2015): Examples of the GPcounts model fit on the expression profiles of two genes (IRF8 and APOE) that are differentially expressed but the expression pattern does not show any evidence of DE at the end of the time range. Cell assignment to different branches as well as the pseudotime of each cell are calculated using the Slingshot algorithm. In both cases, the upper sub-panel shows the GPcounts model fit based on the MAP estimate of the gene-specific differentiating time and the lower sub-panel shows the posterior distribution over the differentiating or branching time point. The bigger markers used in the upper sub-panels to represent the sub-sampled cells that have been used in the inference.

likely that two time courses are truly distinct. On the other hand, if the inferred branching time is at the end of the time, it is very likely that the two samples are similar to each other and are less likely to diverge. Thus a decision can be made over whether or not there is a branching by considering the logged Bayes factor between a model with or without a branching (Boukouvalas *et al.*, 2018). The logged Bayes factor between a model with or without branching is,

$$\begin{aligned} r_g &= \log \frac{p(0 < x_b < x_{\max}|\boldsymbol{y}^t, \boldsymbol{y}^b)}{p(x_b = x_{\max}|\boldsymbol{y}^t, \boldsymbol{y}^b)} \\ &= \log \left[ \frac{1}{N_b} \sum_{x_b = x_{\min}}^{x_b = x_{\max}} p(\boldsymbol{y}^t, \boldsymbol{y}^b | x_b) \right] - \log \left[ p(\boldsymbol{y}^t, \boldsymbol{y}^b | x_{\max}) \right], \end{aligned} \tag{3}$$

where $N_b$ is the number of bins in the histogram approximation to the posterior (see Main paper) and setting $x_b = x_{\max}$ is equivalent to having no branching at all. Equation (3) assumes equal prior probability of having a branching (at any time before $x_{\max}$ with equal probability) or not having a branching. If the height of the posterior at the end of time is greater than the average of the posterior over all earlier times as on the lower sub-panel of Figure 8, the probability of having a branching under the model is less than 0.5. A comprehensive study on the Bayes factor and its applications on different scientific domains are available in Kass and Raftery (1995).

We found $r_g = -1.04$ for gene ERP29. Thus, the logged Bayes factor also suggests a non differentiating scenario. The gene ERP29 is a well known regulator of hematopoiesis, but the data we have are not reflecting branching behaviour, still the model can identify the branching dynamics of this gene. Therefore, the model's efficiency of identifying DE genes relevant to biological plausibilities spreads out a number of different kinds of expression patterns.

### 5 Modelling scRNA-seq pseudotime-series

We apply GPcounts on mouse pancreatic $\alpha$ cell data from scRNA-seq experiments without UMI normalisation (Qiu *et al.*, 2017). We use the pseudotime inference results from Qiu *et al.* (2017) based

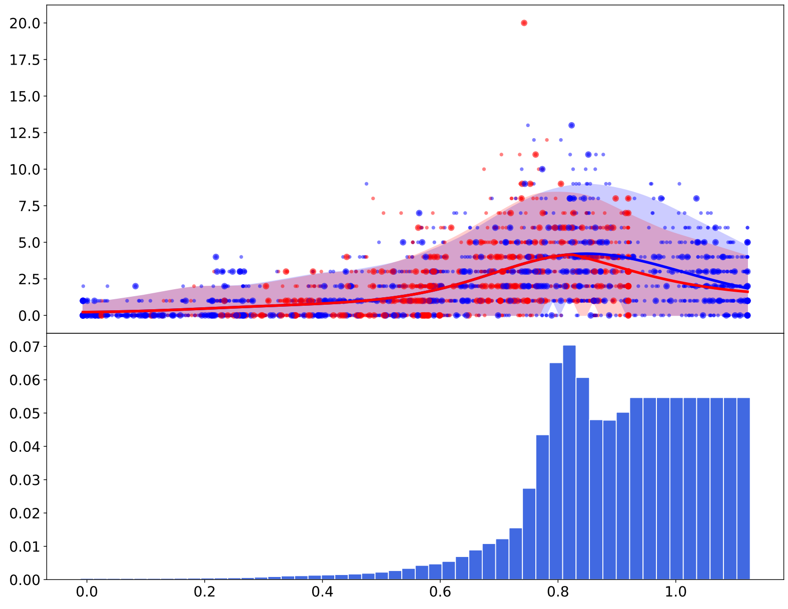


Figure 8: Mouse haematopoietic stem cells (Paul *et al.*, 2015): Example of the GPcounts model fit on gene ERP29. This gene is a well known regulator of hematopoiesis but the expression pattern is showing very little evidence of DE ($r_g = -1.04$). Cell assignment to different branches as well as the pseudotime of each cell are calculated using the Slingshot algorithm. The upper sub-panel shows the GPcounts model fit based on the MAP estimate of the gene-specific differentiating time and the lower sub-panel shows the posterior distribution over the differentiating or branching time point. The bigger markers used in the upper sub-panels to represent the sub-sampled cells that have been used in the inference.

on PCA. Figure 9 and figure 10 shows the inferred trajectory for two genes with many zero count measurements: Fam184b with 86% zero counts and Pde1a with 68% zero counts. From left to right we show the full GP regression fit on the first row with Gaussian, NB and ZINB likelihoods respectively. For Fam184b we see that the Gaussian and NB models do not capture most of the non-zero data within their credible regions. The ZINB model has 21/324 (6.4%) cells above the 95% upper credible region while the NB model has 37/324 (11.4%) suggesting the ZINB model is a little better calibrated. For both genes the Gaussian model is unable to effectively model the high probability region at zero counts, due to the symmetric nature of the distribution. The fits for the NB and ZINB likelihood are very similar for Pde1a with only a subtle difference in the credible regions (4% of cells above 95%) and we find that ZINB and NB give similar fits for most genes.

For scRNA-seq data the number of cells can be very large and computational efficiency becomes important. We show the results of using sparse GP with Gaussian, NB and ZINB likelihoods respectively for Fam184b in figure 9 and for Pde1a in figure 10. In GPcounts, by default we assume the number of inducing points is $M = 5\%(N)$ and we compare the performance of k-mean algorithm (Hensman *et al.*, 2013, 2015) in the second row versus $\epsilon - approximate$ M-DPP algorithm (Burt *et al.*, 2020) in the third row as methods to select the location of the inducing points $\boldsymbol{z}$. We optimize the location of the inducing points selected by k-mean algorithm and fix it for $\epsilon - approximate$ M-DPP algorithm as recommended by methods authors. Using the k-mean algorithm to select the inducing points, improves the computational complexity to be $\mathcal{O}(M^2)$.

For Gaussian likelihood, changing the algorithms does not improve the Spearman correlation scores but for NB likelihood using $\epsilon - approximate$ M-DPP algorithm had improved the correlation by 12 % as shown in figure 11.

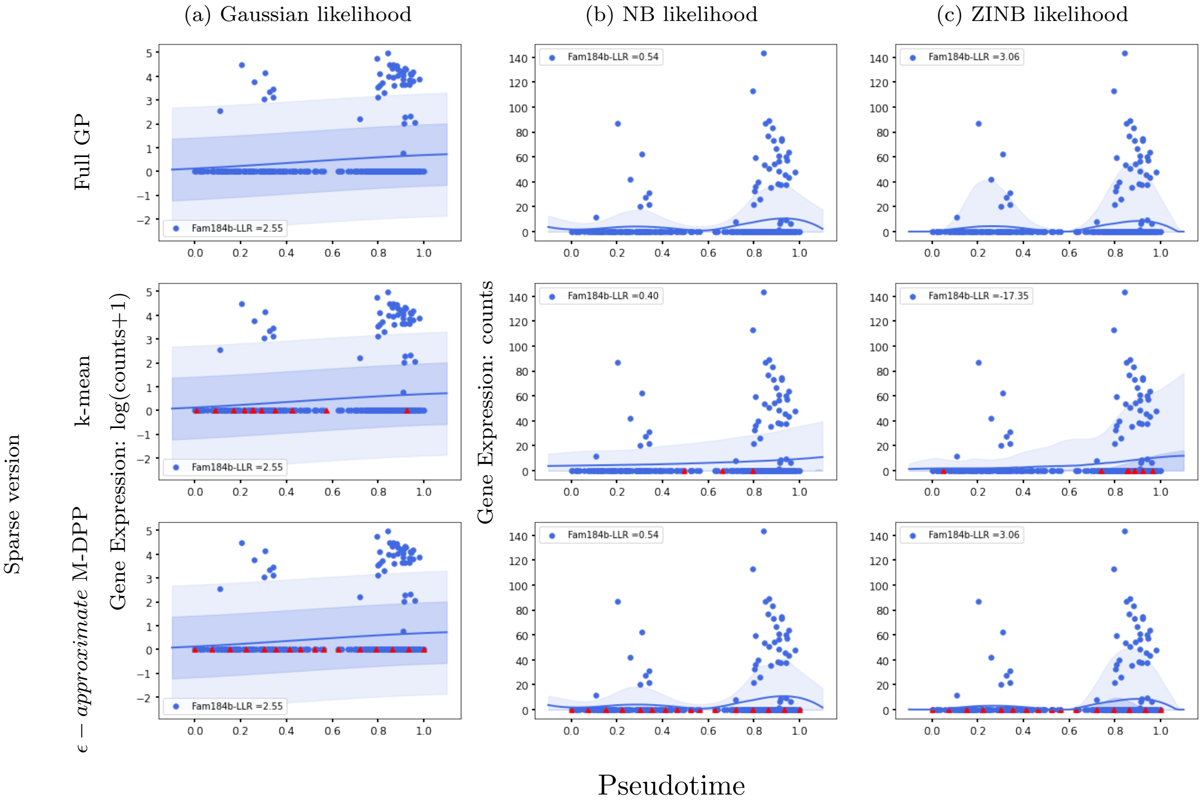


Figure 9: GP models of gene expression against pseudotime (Qiu *et al.*, 2017). We show gene Fam184b with 86% zeros and the inferred mean trajectory and credible regions using (a) Gaussian likelihood, (b) NB likelihood and (c) ZINB likelihood. In the first row we show the fit for a full GP model and compare it with the spare GP fit in the next rows where the numbers of inducing points are $M = 5\%(N)$. We use k-means algorithm in the second row and $\epsilon - approximate$ M-DPP algorithm in the third row to select the locations of the inducing points.

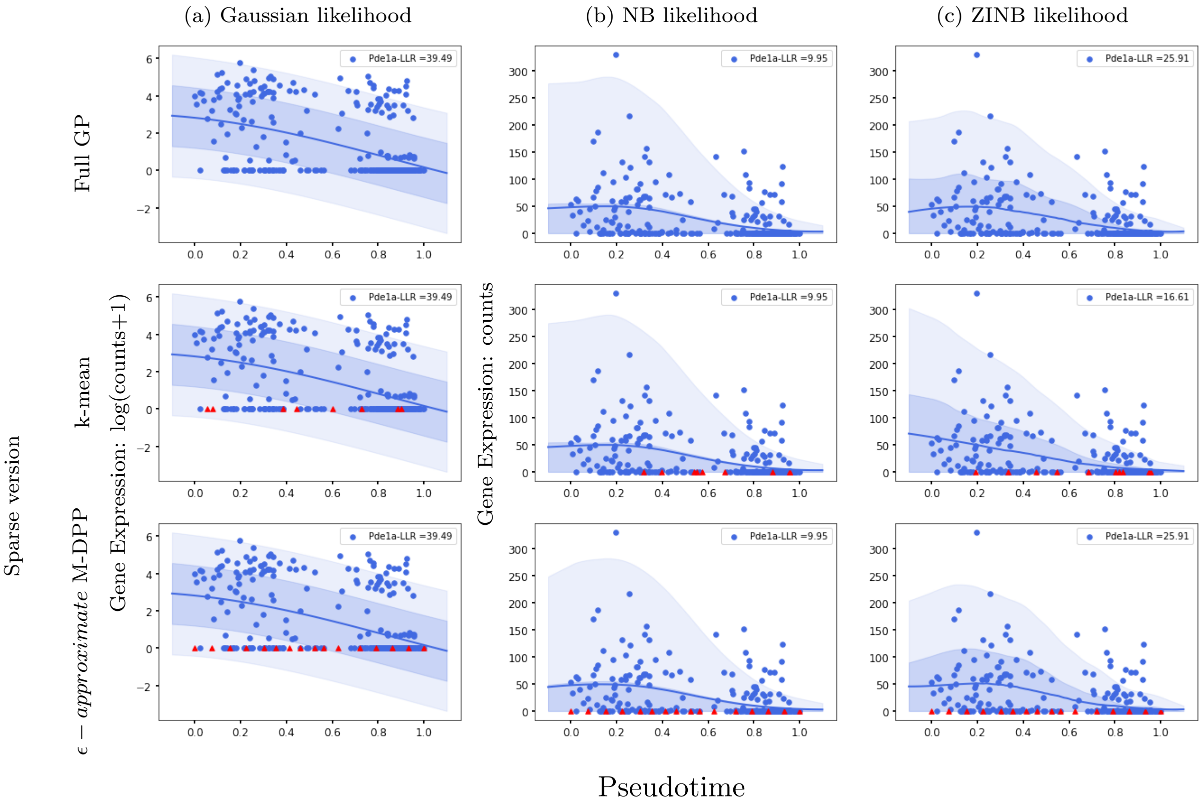


Figure 10: GP models of gene expression against pseudotime (Qiu *et al.*, 2017). We show gene Pde1a with 68% zeros and the inferred mean trajectory and credible regions using (a) Gaussian likelihood, (b) NB likelihood and (c) ZINB likelihood. In the first row we show the fit for a full GP model and compare it with the spare GP fit in the next rows where the numbers of inducing points are $M = 5\%(N)$. We use k-mean algorithm in the second row and $\epsilon - approximate$ M-DPP algorithm in the third row to select the locations of the inducing points.

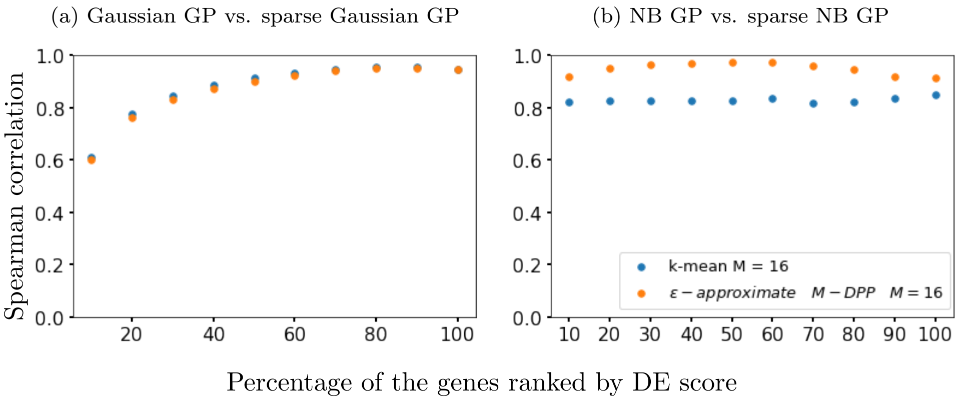


Figure 11: Spearman correlation scores for different percentages of the mouse pancreatic $\alpha$-cell scRNA-seq data (Qiu *et al.*, 2017) ranked by Gaussian GP LLR in (a) and NB GP LLR in (b) . We show likelihood for full Gaussian inference versus different methods to select the number and location of inducing points in sparse GP inference.

### 6 Identification of spatially variable genes

Spatially variable genes are identified based on the comparison between a null hypothesis model and an alternative hypothesis spatial model. The former assumes that there is no difference in gene expression across space, while, the latter, assumes spatial variance. The two models are nested and therefore, their log likelihood ratio ($LLR$) can be used to assess the significance of the alternative hypothesis.

In the original case (Figure 6 of the main document), the $p$-values are computed under the assumption that, the $LLR$ under the null hypothesis follows a $\chi^2$-distribution with one degree of freedom. However, this assumption leads to conservative $p$-values as shown in figure 12a. $P$-values under a permuted null are shown in figure 12b, where the original spots are shuffled as shown in figure 12a. The $p$-value histogram under the permuted null shows a peak near zero, indicating well-behaved $p$-values as well as a larger number of significant genes. This is confirmed in figure 13a and figure 13b, where the number of the spatially variable genes which are called under a permuted null is greater than those under a $\chi^2$-distribution. However, for SpatialDE, this particular random re-arrangement of the spots resulted to even less spatially variable genes (figure 13b). Figure 13c shows the four genes which are only identified as spatially variable by SpatialDE and not by GPcounts, yet these genes are either expressed in one or two locations.

Figure 14(a) shows relative expression profiles of three selected marker genes (Fabp7, Rbfox3 and Eomes) and one house keeper gene (Actb) detected by GPcounts and SPARK as SV. None of these four genes is identified as SV by SpatialDE. Their associated profiles obtained with in-situ hybridization in the Allen Brain Atlas are shown in Figure 14(b).

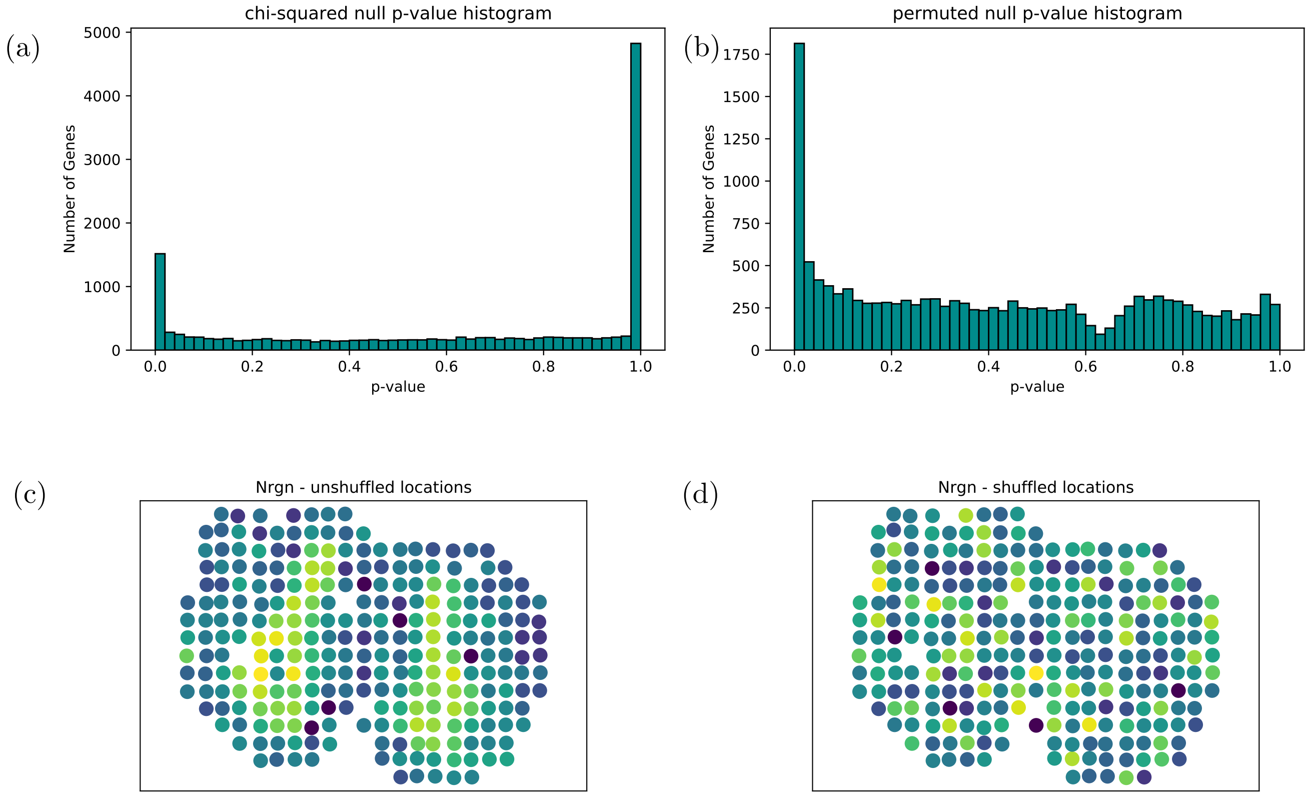


Figure 12: (a) Histogram of $p$-values computed under the assumption that the log-likelihood ratios follow a $\chi^2$-distributed with one degree of freedom null. (b) Histogram of $p$-values computed under a permuted null. (c) Original spatially resolved expression profile of Nrgn (unshuffled locations). (d) Shuffled spatially resolved expression profile of Nrgn. These shuffled locations were used for the permutation test.

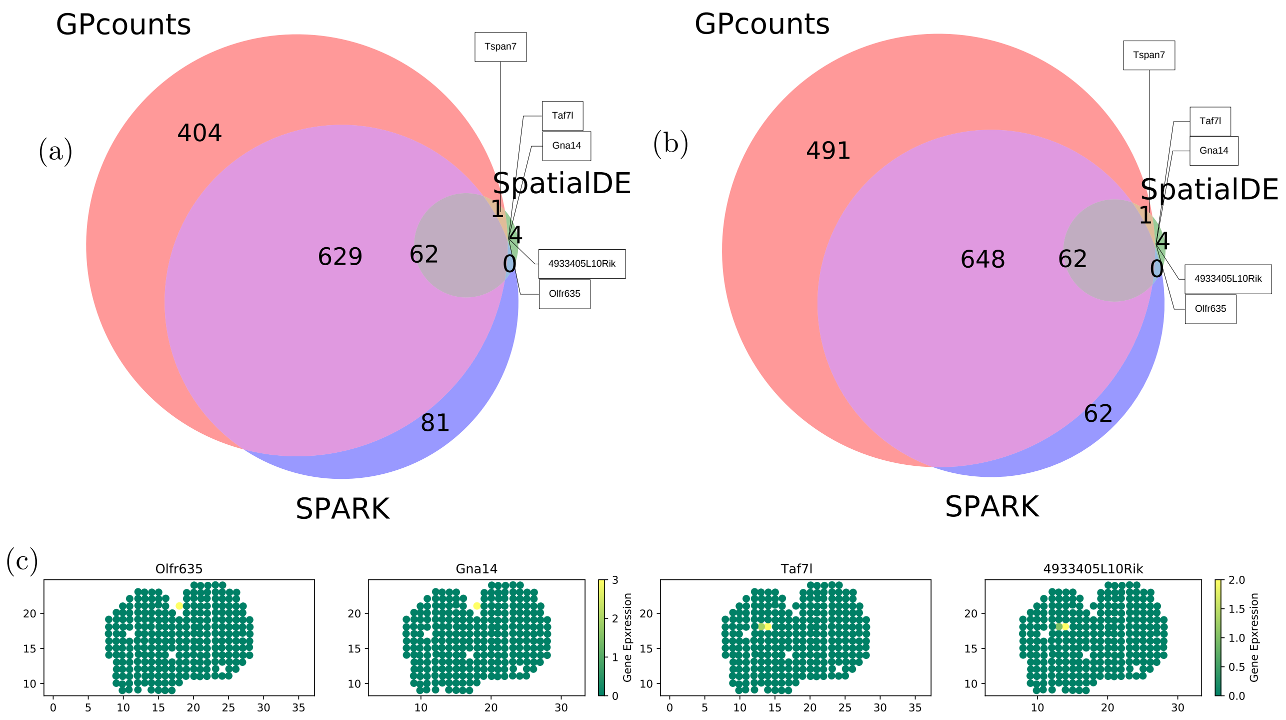


Figure 13: (a) Venn diagram shows the overlap between GPcounts's, SpatialDE's and SPARK's spatially variable genes. The $p$-values of GPcounts and SpatialDE have been computed assuming $\chi^2$-distributed log-likelihood ratios while SPARK computes permuted $p$-values. (b) Venn diagram where, the GPcounts's $p$-values have been computed under a permuted null.(c) Relative spatially resolved expression profiles for the four genes (Gna14, Olfr635, Taf7l and 4933405L10Rik) GPcounts is missing. All four genes show very weak expression profiles.

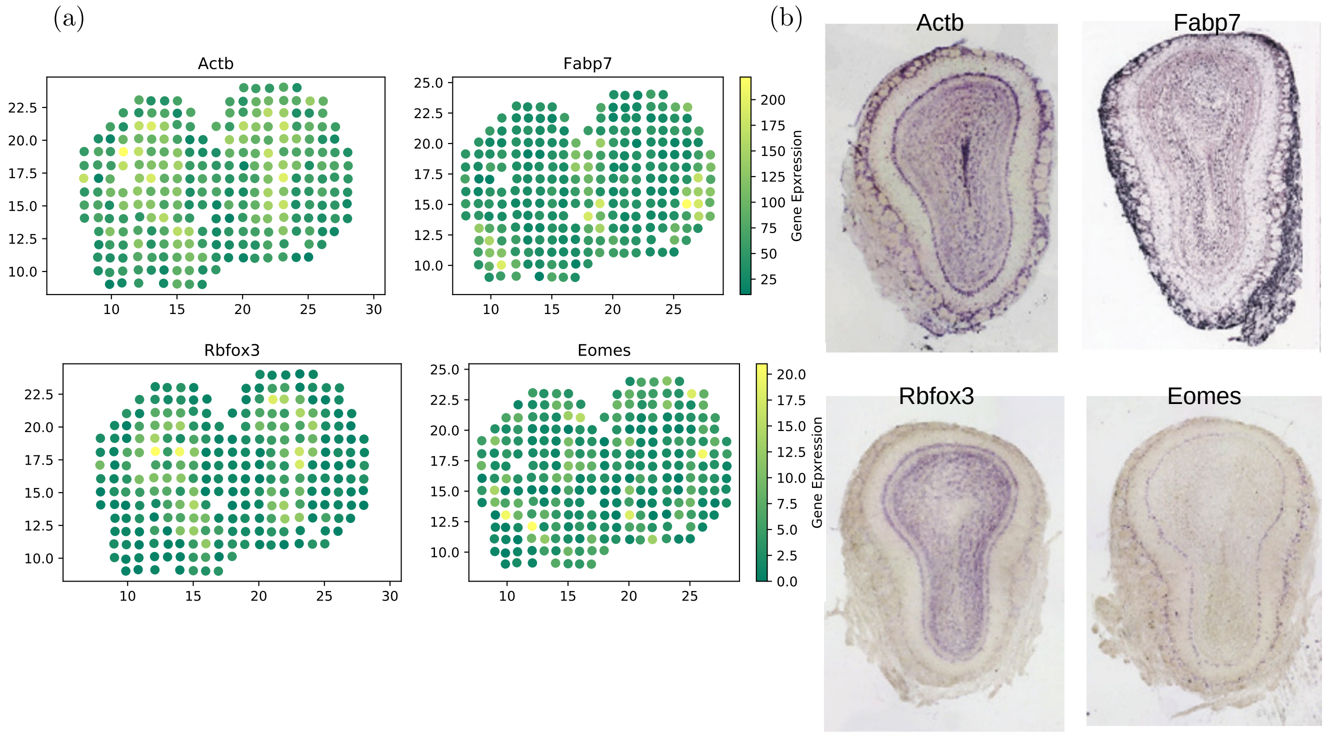


Figure 14: (a) Spatially resolved expression profiles for four selected SV genes. Genes Fabp7, Rbfox3 and Eomes are cluster marker genes while Actb is a house keeper gene. None of these four genes was identified as SV by SpatialDE, while all were identified by GPcounts and SPARK. (b) In-situ hybridization images of the selected SV genes in (a). The images are taken from Ståhl *et al.* (2016).

### References


Andrews, T. S. and Hemberg, M. (2016). Modelling dropouts allows for unbiased identification of marker genes in scRNASeq experiments. *biorxiv*, page 065094.

Boukouvalas, A., Hensman, J., and Rattray, M. (2018). BGP: identifying gene-specific branching dynamics from single-cell data with a branching Gaussian process. *Genome biology*, **19**(1), 65.

Burt, D. R., Rasmussen, C. E., and van der Wilk, M. (2020). Convergence of sparse variational inference in gaussian processes regression. *Journal of Machine Learning Research*, **21**(131), 1–63.

Hensman, J., Fusi, N., and Lawrence, N. D. (2013). Gaussian processes for big data. *arXiv preprint arXiv:1309.6835*.

Hensman, J., Matthews, A., and Ghahramani, Z. (2015). Scalable variational gaussian process classification.

Kass, R. E. and Raftery, A. E. (1995). Bayes factors. *Journal of the american statistical association*, **90**(430), 773–795.

Kurotaki, D., Yamamoto, M., Nishiyama, A., Uno, K., Ban, T., Ichino, M., Sasaki, H., Matsunaga, S., Yoshinari, M., Ryo, A., *et al.* (2014). IRF8 inhibits C/EBP$\alpha$ activity to restrain mononuclear phagocyte progenitors from differentiating into neutrophils. *Nature communications*, **5**, 4978.

Leong, H. S., Dawson, K., Wirth, C., Li, Y., Connolly, Y., Smith, D. L., Wilkinson, C. R., and Miller, C. J. (2014). A global non-coding rna system modulates fission yeast protein levels in response to stress. *Nature communications*, **5**(1), 1–10.

Murphy, A. J., Akhtari, M., Tolani, S., Pagler, T., Bijl, N., Kuo, C.-L., Wang, M., Sanson, M., Abramowicz, S., Welch, C., *et al.* (2011). ApoE regulates hematopoietic stem cell proliferation, monocytosis, and monocyte accumulation in atherosclerotic lesions in mice. *The Journal of clinical investigation*, **121**(10), 4138–4149.

Paul, F., Arkin, Y., Giladi, A., Jaitin, D. A., Kenigsberg, E., Keren-Shaul, H., Winter, D., Lara-Astiaso, D., Gury, M., Weiner, A., *et al.* (2015). Transcriptional heterogeneity and lineage commitment in myeloid progenitors. *Cell*, **163**(7), 1663–1677.

Pierson, E. and Yau, C. (2015). ZIFA: Dimensionality reduction for zero-inflated single-cell gene expression analysis. *Genome biology*, **16**(1), 241.

Qiu, W.-L., Zhang, Y.-W., Feng, Y., Li, L.-C., Yang, L., and Xu, C.-R. (2017). Deciphering pancreatic islet $\beta$ cell and $\alpha$ cell maturation pathways and characteristic features at the single-cell level. *Cell metabolism*, **25**(5), 1194–1205.

Risso, D., Perraudeau, F., Gribkova, S., Dudoit, S., and Vert, J.-P. (2018). A general and flexible method for signal extraction from single-cell rna-seq data. *Nature communications*, **9**(1), 1–17.

Shin, D.-M., Lee, C.-H., and Morse III, H. C. (2011). IRF8 governs expression of genes involved in innate and adaptive immunity in human and mouse germinal center B cells. *PloS one*, **6**(11), e27384.

Ståhl, P. L., Salmén, F., Vickovic, S., Lundmark, A., Navarro, J. F., Magnusson, J., Giacomello, S., Asp, M., Westholm, J. O., Huss, M., Mollbrink, A., Linnarsson, S., Codeluppi, S., Borg, Å., Pontén, F., Costea, P. I., Sahlén, P., Mulder, J., Bergmann, O., Lundeberg, J., and Frisén, J. (2016). Visualization and analysis of gene expression in tissue sections by spatial transcriptomics. *Science*, **353**(6294), 78–82.

Van den Berge, K., De Bezieux, H. R., Street, K., Saelens, W., Cannoodt, R., Saeys, Y., Dudoit, S., and Clement, L. (2020). Trajectory-based differential expression analysis for single-cell sequencing data. *Nature communications*, **11**(1), 1–13.
